# Mating systems and recombination landscape strongly shape genetic diversity and selection in wheat relatives

**DOI:** 10.1101/2023.03.16.532584

**Authors:** Concetta Burgarella, Marie-Fleur Brémaud, Gesa Von Hirschheydt, Veronique Viader, Morgane Ardisson, Sylvain Santoni, Vincent Ranwez, Miguel Navascués, Jacques David, Sylvain Glémin

**Affiliations:** CNRS, Univ Montpellier, ISEM– UMR 5554, 34095 Montpellier, France; AGAP Institut, Univ Montpellier, CIRAD, INRAE, Institut Agro, Montpellier, France; Department of Organismal Biology, Evolutionary Biology Center, Uppsala University, Uppsala, Sweden; Swiss Federal Research Institute WSL, Zürcherstrasse 111 8903 Birmensdorf, Switzerland; UMR CBGP, Univ Montpellier, CIRAD, INRAE, Institut Agro, Montpellier, France; CNRS, Univ. Rennes, ECOBIO – UMR 6553, F-35000 Rennes, France; Department of Ecology and Evolution, Evolutionary Biology Center, Uppsala University, Uppsala, Sweden

**Keywords:** self-fertilization, polymorphism, linked selection, fitness effect of mutations, selfing-syndrome

## Abstract

How and why genetic diversity varies among species is a long-standing question in evolutionary biology. Life history traits have been shown to explain a large part of observed diversity. Among them, mating systems have one of the strongest impacts on genetic diversity, with selfing species usually exhibiting much lower diversity than outcrossing relatives. Theory predicts that a high rate of selfing amplifies selection at linked sites, reducing genetic diversity genome wide, but frequent bottlenecks and rapid population turn-over could also explain low genetic diversity in selfers. However, how linked selection varies with mating systems and whether it is sufficient to explain the observed difference between selfers and outcrossers has never been tested. Here, we used the *Aegilops*/*Triticum* grass species, a group characterized by contrasted mating systems (from obligate out-crossing to high selfing) and marked recombination rate variation across the genome, to quantify the effects of mating system and linked selection on patterns of neutral and selected polymorphism. By analyzing phenotypic and transcriptomic data of 13 species, we show that selfing strongly affects genetic diversity and the efficacy of selection by amplifying the intensity of linked selection genome wide. In particular, signatures of adaptation were only found in the highly recombining regions in outcrossing species. These results bear implications for the evolution of mating systems and more generally for our understanding of the fundamental drivers of genetic diversity.

## Introduction

How and why genetic diversity varies among species is a central and long-standing question in evolutionary biology, dating back from the 1960’s (Ellegren and Galtier, 2016). For neutral variation, patterns of genetic diversity depend on the balance between mutation and genetic drift, characterized by the effective size of a population, *N_e_*, and also on the efficacy of selection for functional regions of the genome. Recently, thanks to the availability of population genomic data in many non-model species, several studies have explored the ecological correlates of diversity levels, usually measured as nucleotide polymorphism, π. These surveys have shown that life history traits (LHTs), especially life-span and reproductive mode, can explain a large part of the observed variation in genetic diversity among species (Romiguier *et al*., 2014; Chen *et al*., 2017; Mackintosh *et al*., 2019; Muyle *et al*., 2021). LHTs may reflect long-term effective population size, which depends on current population size and past fluctuations across generations (e.g. Romiguier *et al*., 2014; Mackintosh *et al*., 2019). *N_e_* can also depend on selection at linked sites, i.e. the hitch-hiking effect of the fixation of beneficial or the removal of deleterious mutations on linked neutral variation (Cutter and Payseur, 2013), which also affects long-term *N_e_* and seems rather pervasive across genomes (Corbett-Detig *et al*., 2015; Mackintosh *et al*., 2019; Chen *et al*., 2020; Buffalo, 2021).

Among LHTs, mating systems deeply affect the genetic and ecological functioning of a species and are predicted to strongly impact both demographic outcomes and the response to selection. Thanks to the ability to produce seeds under limited mate availability, the capacity of autonomous selfing provides reproductive assurance and can be an ecologically successful strategy, allowing colonizing new habitats and increasing species range (Grossenbacher *et al*., 2015), which should be associated with a large census population size. However, being able to reproduce alone implies a much higher demographic stochasticity due to recurrent bottlenecks and colonization-extinction dynamics, which strongly reduces genetic diversity, not only at the population scale but also at the whole species scale (Pannell and Charlesworth, 1999; Ingvarsson, 2002). Moreover, the dynamics of range expansions, which can be associated with the evolution of selfing, can also unintuitively lead to the loss of diversity, especially on the expansion front (Excoffier *et al*., 2009). So, despite possible large species range and census population size, the specific ecology of selfing species may lead to a reduction in *N_e_*. In addition to these demographic effects, selfing also has direct genetic effects that can reduce *N_e_*. Non-independent gamete sampling during mating automatically increases genetic drift and reduces genetic mixing, which generates genome-wide genetic linkage disequilibrium enhancing the effect of linked selection (Agrawal and Hartfield, 2016; Roze, 2016; Hartfield and Bataillon, 2020).

So far, striking differences in genetic diversity have already been observed between outcrossing and selfing relatives (e.g., Hazzouri *et al*., 2013; Slotte *et al*., 2013; Burgarella *et al*., 2015; Teterina et al., 2023) and the underlying causes (linked-selection, demographic instability) have been often discussed and studied from a theoretical point of view (Charlesworth *et al*., 1993; Barrett *et al*., 2014), but, to our knowledge, attempts at a direct quantification with empirical data are recent and only partial (see the comparison between two outcrossing and selfing *Caenorhabditis* species in Teterina *et al*., 2023). Yet, how the intensity of linked selection varies with mating system and whether it can be sufficient to explain the observed difference between outcrossing and selfing species remains to be quantified. Beyond a genomewide reduction in polymorphism and selection efficacy with increasing selfing rates, theory also predicts that genomic patterns across chromosomes should vary with the interaction between recombination and selfing rates. We expect a clear positive relationship between genetic diversity and recombination in outcrossers, but an increasingly flatter relationship in species with increasing selfing rates. We also expect that deleterious mutations should accumulate mainly in lowly recombining regions whereas adaptation should be prevalent in highly recombining regions in outcrossing species, in contrast to selfing species where signatures of high deleterious load and low adaptation should be more evenly distributed along the genome.

Here, we tested these hypotheses by comparing species with a large range of mating systems occurring within a single genus, which allows strong genetic contrast among otherwise similar species, an advocated sampling design (Leffler *et al*., 2012; Cutter and Payseur, 2013). We used the *Aegilops*/*Triticum* grass species as a study system. This group of Mediterranean and Western/Central Asian grasses belongs to the Triticeae tribe (Poaceae) and includes wheat and its wild relatives. The *Aegilops*/*Triticum* genus forms a monophyletic group with 13 diploid and about 17 polyploid species that likely diversified ∼4-7 millions years ago (Huang *et al*., 2002; Marcussen *et al*., 2014; Glémin *et al*., 2019b). All species are characterized by similar life-history traits (wind-pollinated, annual, herbaceous species) and ecology (open landscapes, warm-temperate climate), but present a large diversity of mating systems, spanning from obligate out-crossing to highly selfing species (van Slageren, 1994; Kilian *et al*., 2011) (**Fig. 1**). Triticeae genomes are large, with markedly U-shaped recombination patterns along chromosomes conserved across species: most recombination is located in the distal parts whereas no or very low recombination occurs in their central part (Brazier and Glémin, 2022). Marked differences in both mating systems among species and recombination rate within genomes make the group an ideal model to unravel the role of selection on species genetic diversity.

**Figure 1.**
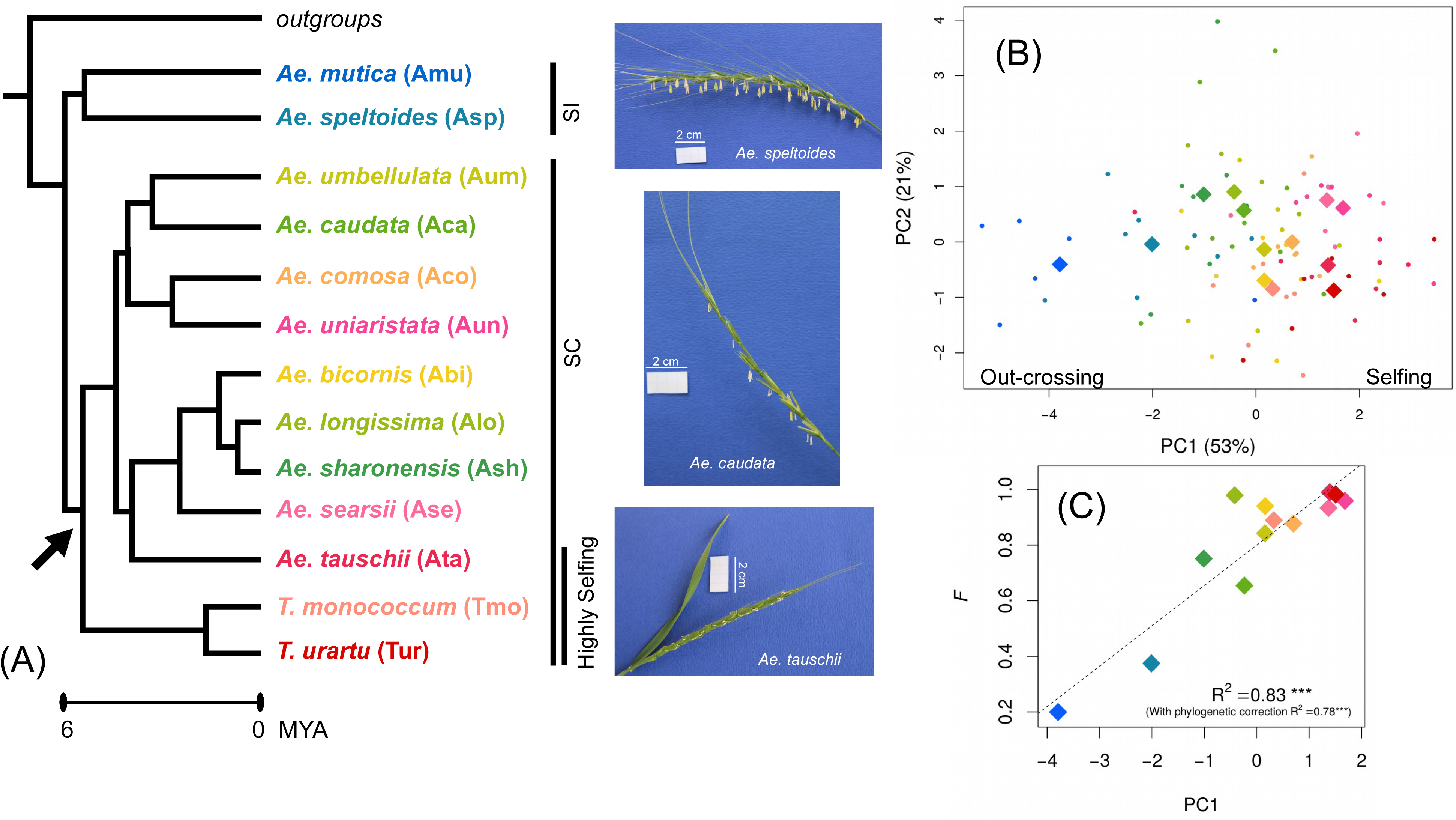
Mating system diversity in the genus *Aegilops*/*Triticum*. (A) Phylogenetic relationships among the 13 diploid species of the genus, following Glémin *et al*. (2019b). Colors have been assigned to represent a gradient of self fertilization: *Ae. speltoides* and *Ae. mutica* have previously been described as self-incompatible (SI, blue tone colors), *Ae. tauschii* and *Triticum* sp. are known to be prevalently self-fertilizing species (pink-purple colors), while the remaining species are self-compatible (SC, yellow-green colors). The arrow indicates when self-compatibility may have appeared, if we assume that it appeared once according to a parsimony approach. Pictures illustrate the phenotypes associated with different mating systems: SI *Ae. speltoides*, highly selfing *Ae. tauschii* and self-compatible *Ae. caudata*. (B) Principal Component Analysis of morphological measures of reproductive organs. PC1 resumes the morphological selfing syndrome. On one side there are the self-incompatible species, which show bigger anther and stigmas, higher male investment, lower spikelet compactness and lower autonomous seed set. On the other side there are predominately selfing species, with opposite states of the traits. (C) Positive correlation between PC1 of reproductive traits and the inbreeding coefficient *F* estimated on the genomic data shows that species with stronger morphological selfing syndrome have also higher *F* values.

Through a comparative population genomic approach, we assessed the expectation that genetic diversity and selection efficacy decreases with higher selfing rate and with lower recombination rate within each species, by controlling for species range that varies among species (**Supplementary Fig. S1**). We then explicitly tested whether the effect of linked selection was stronger in selfing species as predicted by population genetics theory. We found that selfing strongly affects genetic diversity and the efficacy of selection by amplifying the intensity of linked selection genome wide, while species range plays a minor role. We also showed that genomic patterns remarkably matched the gradient of mating systems across species, while models and empirical evidence so far suggested that only extreme mating systems left clear signatures in the genomes (Agrawal and Hardfield 2016, Roze 2016). These results have multiple implications for both the evolution of mating systems and our understanding of the fundamental drivers of genetic diversity.

## Material and methods

### Plant material

We analyzed molecular and morphological data of the 13 extant wild diploid species of the *Aegilops*/*Triticum* genus (Triticeae tribe, Kilian *et al*., 2011) and 3 outgroup species (*Taeniatherum caput-medusae*, *Hordeum spontaneum* and *Secale strictum*) for a total of 98 accessions. We targeted 7-20 individuals from each of five focal species, including the two self-incompatible outcrossing *Ae. speltoides* and *Ae. mutica* and the three predominantly selfing *Ae. tauschii*, *T. urartu* and *T. monococcum*. We included 2-4 accessions from each of the other *Aegilops*/*Triticum* species and 1-3 from each of the outgroups. The accessions were obtained from several international seed banks and donor researchers. The list of accessions per species with their passport information is provided in **Supplementary Table S1**.

### Morphological data

To finely characterize the selfing syndrome of each species, we measured several morphofunctional traits describing the reproductive organs and function (Friedman and Harder, 2005; Escobar *et al*., 2010 and references therein). Around 3-4 grains per accession were sown in January 2014 in a glass house. After emergence, the seedlings were submitted to 4°C for 6 weeks to assure vernalization requirements. Only one seedling per accession was kept after vernalization. The first two spikes of each plant were closed in paper bags to prevent cross fertilization. We collected the two bagged spikes, one open mature spike, mature anthers, stigmas and ovaries for measurements. Anthers, stigmas and ovaries were preserved in Carnoy fixative solution. For each of 5-7 accessions per *Aegilops*/*Triticum* species and 1-3 accessions per outgroup species, we measured a mature spike (length, spikelet number, grain number), three spikelets per spike (spikelet length) and three flowers from one spikelet (length of palea and lemma). All flowers of the three spikelets were classified as fertile (if the presence of grain was observed), female (only stigma observed), male (anthers observed), or sterile. For each accession, we measured 6 anthers (length and width) and 3 stigmas and ovaries (length of each organ). For anthers and stigma and ovaries, each organ was measured five times and the mean value of these replicates was used in further analysis. Measures were manually recorded on millimeter paper or taken on photographs with the software analySIS (Soft Imaging System GmbH 2002; see **Supplementary Fig. S12** for an example).

Missing data (31%) on the directly observed measures were imputed with the missMDA package (Josse and Husson, 2016) under the R environment (R Core Team, 2018). Parameters were set by default and the optimal number of components retained for imputation were estimated with the cross-validation method (ncp=3). Raw and imputed measures are provided in **Supplementary tables S2** and **S3**, respectively and the list of measured traits is provided in **Table S4**.

Additional variables were calculated on imputed measures as follows. The mean values of anther and stigma dimensions were calculated per accession and standardized by dividing by the flower length. Following Escobar *et al*. (2010), the autonomous seed set was estimated as self-fertilised_seed_number/(self-fertilized_spikelet_number*number_fertile_flower/spikelet), corresponding to the number of seeds per fertile flower. Spikelet compactness was calculated as the ratio (mean flower length* number_fertile_flower)/mean spikelet length. Male investment was calculated as the ratio of the mean anther length and the mean ovary length. We used these additional variables to summarize the selfing syndrome with a synthetic measure corresponding to the first axis of a Principal Component Analysis (PCA) (**Supplementary Table S3** and **S4**). The PCA was performed with the ade4 package (Dray and Dufour, 2007) under the R environment.

### Species range

We expect that species with bigger census sizes also harbor higher genetic diversity, a relationship that could mask or interact with the effect of the mating system. To control for this potential effect, we used species range as a proxy for census size, since, to our knowledge, there are no direct estimates of census size for wild *Aegilops*/*Triticum* species. To estimate species range, we retrieved occurrence data from the Global Biodiversity Information Facility (http://www.gbif.org) for each species. We manually cleaned the data set to remove single occurrences outside species range, which can be due either to identification errors or recent introductions. Cleaned data were mapped on the world map (focusing on western Eurasia and North Africa) on which we applied a grid with cell size of one decimal degree square (∼10,000 km^2^). We estimated species range as the number of cells occupied by a species time 10,000 km^2^ (**Supplementary Figure S1 and Table S5**).

### Sequencing

We added 48 new sequences to the dataset used for the phylogenomic analysis of Glémin *et al*. (2019b) and Clément *et al*. (2017), for a total of 98 sequences for 13 *Aegilops*/*Triticum* species (n=2-21) and 3 outgroup species (n=1-3).

We performed full transcriptome sequencing following the procedure described in Sarah *et al*. (2017) and Glémin *et al*. (2019b). Briefly, RNAs were extracted and prepared separately for leaves and inflorescence tissues, and mixed subsequently in 20% and 80% proportions, respectively. RNA was extracted using a Spectrum Plant Total RNA kit (Sigma-Aldrich, USA) with a DNAse treatment. RNA concentration was measured with two methods, a NanoDrop ND-1000 Spectrophotometer and the Quant-iT™ RiboGreen® (Invitrogen, USA) protocol. RNA quality was assessed on the RNA 6000 Pico chip on a Bioanalyzer 2100 (Agilent Technologies, USA). Following the Illumina TruSeq mRNA protocol, we kept samples with an RNA Integrity Number (RIN) value greater than eight. Libraries were prepared with a modified protocol of the TruSeq Stranded mRNA Library Prep Kit (Illumina, USA) to obtain library fragments of 250-300 bp. Modification details and amplification conditions are available in Glémin *et al*. (2019b). After verifying and quantifying each indexed cDNA library using a DNA 100 Chip on a Bioanalyzer 2100, pooled libraries were made of twelve, equally represented, genotypes. Each final pooled library was quantified by qPCR with the KAPA Library Quantification Kit (KAPA Biosystems, USA) and sequenced using the Illumina paired-end protocol on a HiSeq3000 sequencer by the Get-PlaGe core facility (GenoToul platform, INRA Toulouse, France http://www.genotoul.fr).

### Transcriptome assembly, mapping and genotype calling

Reads cleaning and assembly were performed with the pipeline described in Sarah *et al*. (2017). Adapters were removed with cutadapt (Martin, 2011). Reads were trimmed at the end, removing sequences with low quality scores (parameter –q 20), and we retained only reads with a minimum length of 35 bp and a mean quality higher than 30. Orphan reads were then discarded using a homemade script. Retained reads were assembled with ABySS (Simpson *et al*., 2009), using the paired-end option with a kmer value of 60, followed by one step of Cap3 (Huang and Madan, 1999) run with the default parameters, 40 bases of overlap and 90% percentage of identity. To predict the CDS embedded in our contigs, we used the *prot4est* program (Wasmuth and Blaxter, 2004). We provided three gene datasets: the output of a *Rapsearch* (Ye *et al*., 2011) similarity analysis, *Oryza* matrix model for de-novo based predictions and the codon usage bias observed in *T. monococcum*. We run Rapsearch to identify protein sequences similar to our contigs in either plant species of Uniprot swissprot (http://www.uniprot.org) or in the Monocotyledon species of greenphyl (http://www.greenphyl.org/cgi-bin/index.cgi). For the individual used as mapping reference within each species (see below), we discarded predicted CDS with less than 250bp.

For each species, mapping was done following Sarah *et al*. (2017), except for the use of the bwa (Li and Durbin, 2009) option –mem (instead of –aln) more adapted for reads of 100 bp. Reads were mapped on the sequences of the individual with the highest coverage or with the highest number of annotated contigs. The list of samples used as reference sequences and the total number of contigs per reference is given in **Supplementary Table S5**.

For each individual, diploid genotypes were called with reads2snps v. 2.0.64 (Tsagkogeorga *et al*., 2012; Gayral *et al*., 2013) (available at https://kimura.univ-montp2.fr/PopPhyl/index.php?section=tools). This tool is specifically designed to analyze transcriptome data for population genomics of non-model species. The method first calculates the posterior probability of each possible genotype in the maximum-likelihood framework, after estimating the sequencing error rate. Genotypes supported with probability higher than a given threshold (here 0.95) are retained, otherwise missing data are called. We required a minimum coverage of 10X per position and per individual to call a genotype. SNPs are then filtered for possible hidden paralogs (duplicated genes) using a likelihood ratio test based on explicit modeling of paralogy (“paraclean” option embedded in the reads2snps software, (Gayral *et al*., 2013)). First, genotype and SNPs were called assuming panmixia (heterozygote deficiency *F* = 0), and *F* was estimated on the retained SNPs. As we have species-wide samples, *F* is equivalent to a *F*_IT_ and mainly corresponds to *F*_IS_ for selfing species and to *F*_ST_ for outcrossing ones. As the assumed expected heterozygosity can affect genotype calling and paralog filtering, reads2snps were run a second time for each species using the *F* estimated after the first step. For the outgroup species with sample size n=1 (*T. caput-medusae*) we kept the initial genotype calling and filtering procedure.

Open-reading frames (ORFs) were predicted using the program ORF_extractor.pl (available at https://kimura.univ-montp2.fr/PopPhyl/index.php?section=tools). Gene length and number of SNPs in the final data set per species is provided in **Supplementary Table S5**.

Orthologous pairs of ORFs, hereafter called genes, from the 5 focal and one outgroup species (*T. caput-medusae*) were identified using reciprocal best hits on BLASTn results, a hit being considered as valid when *e* value was below e-50. Outgroup sequences were added to within-focal species alignments using MACSE v. 1.2 (Ranwez *et al*., 2011), a program dedicated to the alignment of coding sequences and the detection of frameshifts. Genes were only retained if no frameshift was identified by MACSE, and if the predicted ORF in the focal species was longer than 100 codons.

### Chromosome patterns and recombination map

We wanted to analyze polymorphism patterns across chromosomes and as a function of recombination rates. Unfortunately, there is neither reference genome nor genetic map available for every species. Among the high-quality recombination maps available (see Brazier and Glémin 2022), we first used the recombination map of *Hordeum vulgare* as a reference for all species. The synteny is well conserved at the scale of Triticeae (Mayer *et al*., 2011; but see Parisod and Badaeva, 2020) and, as *H. vulgare* is an outgroup, there should not be a specific bias for one species or another. For comparison, we also used the three constitutive genomes AA, BB and DD of the hexaploid wheat, *Trticum aestivum*, which correspond to the three main lineages in the *Aegilops*/*Triticum* phylogeny (Glémin et al. 2019 and **Fig. 1**). These genomes are closer to the focal species but the phylogenetic distance depends on the genome and the species.

To build the recombination maps, we built genetic versus physical distance maps (Marey’s maps). For the barley genome, we used the genetic SNP markers from Comadran *et al*. (2012) which was initially mapped on version 082214v1 of the barley genome. We thus used the coordinate correspondence between this first version and the new reference genome assembly (Hv_IBSC_PGSB_v21) to locate the SNP markers on this reference genome. After visual inspection of aberrant markers, we kept a total of 3590 markers (on average ∼513 markers per chromosome). Recombination rates were computed with the *MareyMap* R package (Rezvoy *et al*., 2007) by fitting a loess function with a second-degree polynomial on sliding windows containing 20% of the markers of a chromosome. This led to a rather smooth recombination map, which is sufficient for our purpose of capturing large-scale patterns and reducing noise. For the bread wheat genome, we used the Marey maps built in Brazier and Glémin (2022). Genetic distances were then interpolated between markers using the fitted function so that a genetic distance could be attributed to each annotated gene of the *Hordeum* genome. For each gene we computed the local recombination rate by taking the local derivative of the fitted loess function as implemented in MareyMap (recombination maps are provided in **Supplementary Fig. S13**).

For each assembled transcript of each focal species, we searched for its orthologous sequence in the *H. vulgare* genome (H) using reciprocal best blast and retaining pairs when *e* value was below e-50. Then the two sequences were aligned with MACSE v.1.2 and the synonymous divergence, *D*_S_, was computed using *codeml* from the PAML software (Yang, 2007). The *D*_S_ distribution was clearly bimodal for all species, and gene pairs showing too high divergence (*D*_S_ > 0.35) were discarded as they likely corresponded to paralogues. For each transcript in each focal species, we attributed the same genetic distance and local recombination rate as its ortholog in *H. vulgare*. We applied the same procedure for the three *T. aestivum* subgenomes (A, B and D) separately, except that we did not filtered on *D*_S_ that cannot be used as an homogeneous criterium for all species as the distance depends on the subgenome and the focal species, contrary to *Hordeum*, which as the same expected distance with all focal species.

For each focal species, to assess the similarity among genomes, we counted how many genes had an orthologue on the same chromosome of the four reference genomes (H, A, B and D) and whether it corresponded to the same category of recombination (low: below the median, or high: above the median).

### Sequence polymorphism analysis

Polymorphism and divergence statistics were calculated with dNdSpNpS v.3. (available at https://kimura.univ-montp2.fr/PopPhyl/index.php?section=tools) that rely on the Bio++ libraries (Guéguen *et al*., 2013). Further filters were applied to the data sets. Positions at which a genotype could be called in less than five individuals for species with sample size n ≥ 5 and in less than n/2 for species with n < 5 were discarded. Genes with less than 10 codons were discarded. For each gene, the following statistics were calculated: per-site synonymous (π_S_) and nonsynonymous (π_N_) mean pairwise nucleotide diversity, heterozygote deficiency (*F*), number of synonymous (S_S_) and nonsynonymous (S_N_) segregating sites, number of synonymous (*D*_S_) and nonsynonymous (*D*_N_) fixed differences between focal and outgroup species. These statistics were computed from complete, biallelic sites only, i.e., sites showing no missing data after alignment cleaning, and no more than two distinct states. For each species, statistics were averaged across genes weighting by the number of complete sites per gene, thus giving equal weight to every SNP. For π_N_/π_S_ and *D*_N_/*D*_S_, we first computed the averages of π_N_, π_S_, *D*_N_, and *D*_S_ and subsequently the ratios of averages. Confidence intervals were obtained by 10,000 bootstraps over genes. For the focal selfing species *Ae. tauschii*, *T. monococcum* and *T. urartu*, all statistics were calculated on n/2 alleles, by randomly drawing one haploid sequence per gene and individual (**Supplementary Table S5**).

### Fit of a linked-selection model

To go further, we fitted a linked-selection model, following Corbett-Detig *et al*. (2015) and Elyashiv *et al*. (2016) but including the effect of partial selfing. To simplify the model, and because we did not have information about substitutions across the genome, we only considered background selection.

The genome was split into genomic windows. For each region where *π_S_* (*i*) has been estimated on *n_i_* positions, we assumed that *n_i_* x *π_S_* (*i*) followed a binomial distribution with parameter *n_i_* and *p_i_* given by:

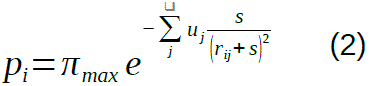

where *s* is the mean selection coefficient against deleterious mutations, *r_ij_* represents the probability of recombination between the focal region *i* and any other region of the genome, *j* containing *L_j_* coding positions so that *u_j_ = u L_j_*, and where *u* is the rate of deleterious mutations. To improve the fit to the data we considered a distribution of fitness effects of mutations. We used a simple discrete distribution with three categories characterized by their mutation rates and selection coefficients: *u_1_*, *u_2_*, *u_3_*, and *s_1_*, *s_2_*, *s_3_*. Equation (2) can thus generalized as:

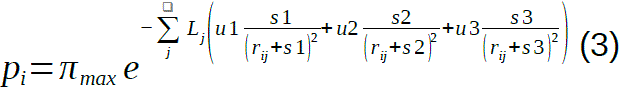

Note that the *s* values correspond in fact to (*h + F – hF*)*s* where *h* is the dominance coefficient. However, because our aim is not to estimate and compare deleterious mutation parameters we did not fit *h* and *s* separately. *r*_ij_ is obtained from the genetic distance, *d*_ij_, using Haldane’s mapping function: *r*_ij_ = (1 – exp(–2*d*_ij_))/2. To take partial selfing into account, we rescaled *r*_ij_ by *r*_ij_(1 – *F*) (see Nordborg, 2000). This rescaling is correct only for low recombination rates and a more accurate expression of background selection was obtained by Roze (2016). However, the simple rescaling provides a good approximation and is much simpler to handle than the full expression (Roze, 2016). Note that we took the sum on all genomic regions on the same chromosome of the focal region but also on other chromosomes (so with *r_ij_* = ½). This is especially important under partial selfing as selection on one chromosome can also affect the other chromosomes. Under outcrossing it boils down to the effect of the additive variance in fitness that reduces effective population size (Roze, 2016).

We had seven parameters to estimate: *u_1_*, *u_2_*, *u_3_*, *s_1_*, *s_2_*, *s_3_* and π*_max_*. Because estimates of *F* were not very precise, we run the model by letting *F* free and being estimated jointly with the other parameters (so eight parameters in total). The log likelihood function was optimized with the *optim* function in *R* using the “L-BFGS-B” method and the constraint: *u_1_* > *u_2_* > *u_3_*.

Recombination is very heterogeneous along chromosomes (U-shaped) and large central regions are strongly linked. Instead of splitting the genome into regions of equal physical size (in Mb) we split it into regions of equal genetic size (in cM) based on the Marey map interpolation. Genomic regions on the telomeric parts of the chromosomes were thus shorter than centromeric regions, where recombination is very low. We chose a window size of 1 cM and we discarded regions with less than 300 bp to avoid too noisy data. To obtain confidence intervals on parameter estimations we bootstrapped data 100 times and rerun the model.

### Linear models and phylogenetic correction

We looked at the relation of polymorphism (π_S_, π_N_/π_S_ and*π_max_*) and *F* with mating system (PC1 of reproductive morphology) and species geographical range using simple unweighted linear regressions of the form *y* ∼ *x* (*F* ∼ PC1, log(π_S_) ∼ PC1, π_N_/π_S_ ∼ PC1, log(π_S_) ∼ range, π_N_/π_S_ ∼ range, π_N_/π_S_ ∼ *F*). To evaluate the joint effects of mating system and species range on polymorphism (π_S_ and π_N_/π_S_), we performed multiple linear regressions in the form log(π_S_) ∼ mating_system + species range and π_N_*/*π_S_ ∼ mating_system + species range. We then represented the residuals, after removing the effect of the mating system, as a function of species range. All linear models were run with the function *lm* under the R environment.

We also applied a correction to take into account the phylogenetic relationships among species using the ultrametric tree retrieved from Glémin *et al*. (2019). For this, we computed the phylogenetically independent contrasts using the method of Felsenstein (1985) with the package *ape* version 5.6-2 (Paradis and Schliep, 2019). The function *pic* was applied to the *y* and *x* vectors with default parameters, then the *lm* analysis was repeated with the contrast values obtained instead of the raw values.

### Estimation of the distribution of fitness effect of mutations

In the two self-incompatible species, *Ae. mutica* and *Ae. speltoides*, and in two highly selfing species, *Ae. tauschii* and *T. urartu*, we estimated the distribution of fitness effect of mutations using the *PolyDFE* method (Tataru *et al*., 2017; Tataru and Bataillon, 2019). In brief, this method used the unfolded site frequency spectrum (uSFS) for both synonymous and non-synonymous mutations to fit the distribution of fitness effect (DFE) of mutations modelled by the mix of a gamma distribution for deleterious mutations and an exponential distribution for beneficial mutations. Demography is taken into account by adding and fitting noise parameters that distort uSFS from the equilibrium expectation following Eyre-Walker *et al*. (2006). Because uSFS are sensitive to polarization errors, which can give spurious signatures of beneficial mutations, a probability of mis-polarization is also added and fitted in the model. This yields a set of four related models: with and without beneficial mutations, and with and without polarization errors. Instead of choosing the best model to estimate parameters, we run all four and used a model averaging procedure (as in Muyle *et al*., 2021): each parameter estimate was averaged using Akaike weights 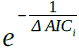, with *Δ AIC_i_*= *AIC_i_ − AIC_min_*where *AIC_min_* is the AIC of the best model. Confidence intervals were obtained by bootstrapping SNPs 1000 times.

For the four species, uSFS were polarized using *Taeniatherum caput-medusae* as an outgroup. We performed three analyses: on the whole set of SNPs or by splitting the dataset into two subsets: SNPs from genes in the high or low recombining regions. GC-biased gene conversion (gBGC), a recombination associated processes mimicking selection in favor of G and C nucleotides, is known to be active in grasses (Rodgers-Melnick *et al*., 2016; Clément *et al*., 2017) and could lead to spurious signatures of positive selection in highly recombining regions. To test for the possible effect of gBGC, we also re-run the analyses on three categories of SNPs: AT→GC, GC→AT, and G←→C + A←→T.

### Simulations

To understand how different selfing rates affect our ability to detect the effects of linked selection on polymorphism landscape and distribution of fitness effects, we run forward time individual-based evolutionary simulations in SLiM v.3.3 (Haller and Messer 2019). We simulated a population of *N* = 10,000 individuals with five different selfing rates: 0, 0.5, 0.9, 0.99 and 1. We considered a genome of three Mb with a single chromosome composed of 1000 genes of 1000 bp separated by intergenic regions of 2000 bp. Recombination decreased exponentially from 60 cM/Mb at the tips to 6 cM/Mb in the center of the chromosome, corresponding to a total genetic of 3.24 Morgan (so an average of three crossovers per chromosome per meiosis, which is in the range of one to three/four that is observed in plants, Brazier and Glémin 2022). We assumed a mutation rate of 10^-6^, with ⅔ of mutations being neutral, corresponding to an expected genetic diversity of 4*Nu* = 0.027, of the order of magnitude of what we observed in the outcrossing species, and corresponding to an average *r*/*u* close to one. The other third of mutations were considered deleterious with a dominance level of *h* = 0.25 and deleterious effects in homozygotes drawn in a gamma distribution with mean = 0.01 and shape = 0.5. This corresponds to a genomic deleterious mutation rate of *U* = 0.33. After a burn-in period of 10*N* generations we recorded the genome sequence of 15 individuals. We run ten replicates for each selfing rate.

We used the simulated data to assess the effect of selfing and linked selection on the estimation of the DFE. Importantly, polyDFE estimates the shape and the population-scaled mean of the DFE: *S* = 4*N_e_*(*h* + *F* – *hF*)*s*. However, *N_e_* is not set nor fixed in the model, contrary to *h*, *s* and *F*, but depends on the intensity of linked selection. We thus used the observed π*_S_* divided by 4*u* to get the predicted *N_e_*, hence the predicted *S*.

## Results

### Mating system widely varies in *Aegilops*/*Triticum* genus

We analyzed phenotypic and transcriptomic diversity in 98 accessions from the 13 diploid *Aegilops*/*Triticum* species and three close outgroup species *Taenatherium caput-medusae*, *Hordeum vulgare* and *Secale vavilovii* (**Fig. 1, Table S1** and **Fig. S1**). Individuals were sampled over the whole geographic range to assess genetic diversity at the species scale. For some species the mating system was already well known, including the self-incompatible *Ae. speltoides* and *Ae. mutica* and the highly selfing *T. urartu*, *T. monococcum* and *Ae. tauschii* (Dvořák *et al*., 1998; Escobar *et al*., 2010), but for others it was poorly documented (Kilian *et al*., 2011). We thus characterized the mating system of each species by quantifying six floral and reproductive traits, including the size of female and male reproductive organs (anthers, stigmas), male investment, spikelet compactness, and the autonomous seed set (Escobar *et al*., 2010) (**Supplementary Tables S2, S3 and S4**). Building on previous work (Escobar *et al*., 2010), we considered these traits as indicative of the selfing syndrome, i.e. the specific changes in flower morphology and function that are expected to occur following the evolution of self-fertilization, especially for anemophilous species (Escobar *et al*., 2010; Sicard and Lenhard, 2011). The autonomous seed set provided a verification that self-incompatible species *Ae. mutica* and *Ae. speltoides* produced almost no seeds under imposed self-fertilization in the greenhouse (bagged spikes), while all the other species were able to produce seeds (**Supplementary Fig. S2**). All the other traits were significantly negatively (anther and stigma size, male investment) or positively (spikelet compactness) correlated with the autonomous seeds set (**Supplementary Fig. S3**), indicating that the selected traits are good indicators of the mating system of each species.

We summarized this reproductive morpho-functional diversity with a multivariate approach, principal component analysis (PCA). The PCA first axis reflected the differences in mating systems within the *Aegilops*/*Triticum* genus (**Fig. 1B** and **Supplementary Fig. S4**). On one extreme of PC1 there were self-incompatible *Ae. mutica* and *Ae. speltoides*, which showed bigger anthers and stigmas, higher male/female investment, lower spikelet compactness and lower (null) autonomous seed set. On the other extreme there were the predominantly selfing *Ae. tauschii*, *T. urartu* and *T. monococcum*, with opposite states of the traits (**Fig. 1A and 1B**). The other *Aegilops* species showed intermediate values of the multi-trait statistic. This result still held when outgroups were included in the analysis (**Supplementary Fig. S5**).

Species morphology measured by PC1 explained well the genome-wide estimate of inbreeding coefficient, *F,* calculated on the whole transcriptome dataset (R^2^=0.83, p-value=1.68e-05, **Fig. 1C**), confirming that our phenotypic data represented a good proxy of the mating system. Thus, in the following analyses, we used PC1 to describe the selfing syndrome, which avoids using genomic data for both characterizing the mating systems and their genomic consequences.

These findings are in agreement with previous knowledge (Dvořák *et al*., 1998; Escobar *et al*., 2010) and allowed us to characterize the mating system for the species that lacked outcrossing rate estimations. They also showed that the effects of selfing might be gradual, with species exhibiting mixed mating strategies described by intermediate values of *F* and of the phenotypic selfing syndrome. Mapping mating systems on the phylogeny suggested that self-incompatibility was likely ancestral and may have broken only once as all species are self-compatible, except the two external ones (*Ae. mutica* and *Ae. speltoides*). However, several breakdowns of self-incompatibility cannot be excluded. For example, high selfing could have evolved four times independently, in the branches leading to *Ae. tauschii*, *Ae. searsii*, *Ae. uniaristata* and *Triticum* species (**Fig. 1A**).

### Polymorphism strongly correlates with mating systems

Genetic diversity was estimated for each species from whole transcriptome sequencing data. Sequences of 48 accessions were generated and *de novo* assembled in this study and were added to the datasets of Glémin *et al*. (2019b) and Clément *et al*. (2017) (**Supplementary Table S1**). Between 19,518 and 28,834 coding sequences were obtained per species. After filtering, genotype calling was performed on a number of contigs varying from 7,083 (*Ae. searsii*) to 21,706 (*T. caput-medusae*) (**Supplementary Table S5**).

Selfing is expected to reduce neutral genetic diversity (Pollak, 1987; Schoen and Brown, 1991; Jarne, 1995; Nordborg, 2000; Ingvarsson, 2002), here estimated as synonymous polymorphism, π_S_. Across species, π_S_ varied more than one order of magnitude, from 0.0011 for the self-compatible *Ae. searsii* to 0.02 for the self-incompatible *Ae. speltoides*. According to expectations, genetic diversity decreases with increasing selfing rate. Neutral genetic diversity, π_S_ and PC1 were significantly correlated across species (R^2^ = 0.75, p-value = 0.00014), indicating a gradient in which stronger selfing syndrome corresponds to lower genome-wide neutral diversity (**Fig. 2A**). The correlation was still significant after phylogenetic control (R^2^ = 0.67, p-value = 0.00111). Interestingly, this relationship is more or less log-linear (**Fig. 2A**) with the main difference being observed between the two self-incompatible (*Ae. speltoides* and *Ae. mutica*) and the self-compatible species (all the others).

**Figure 2.**
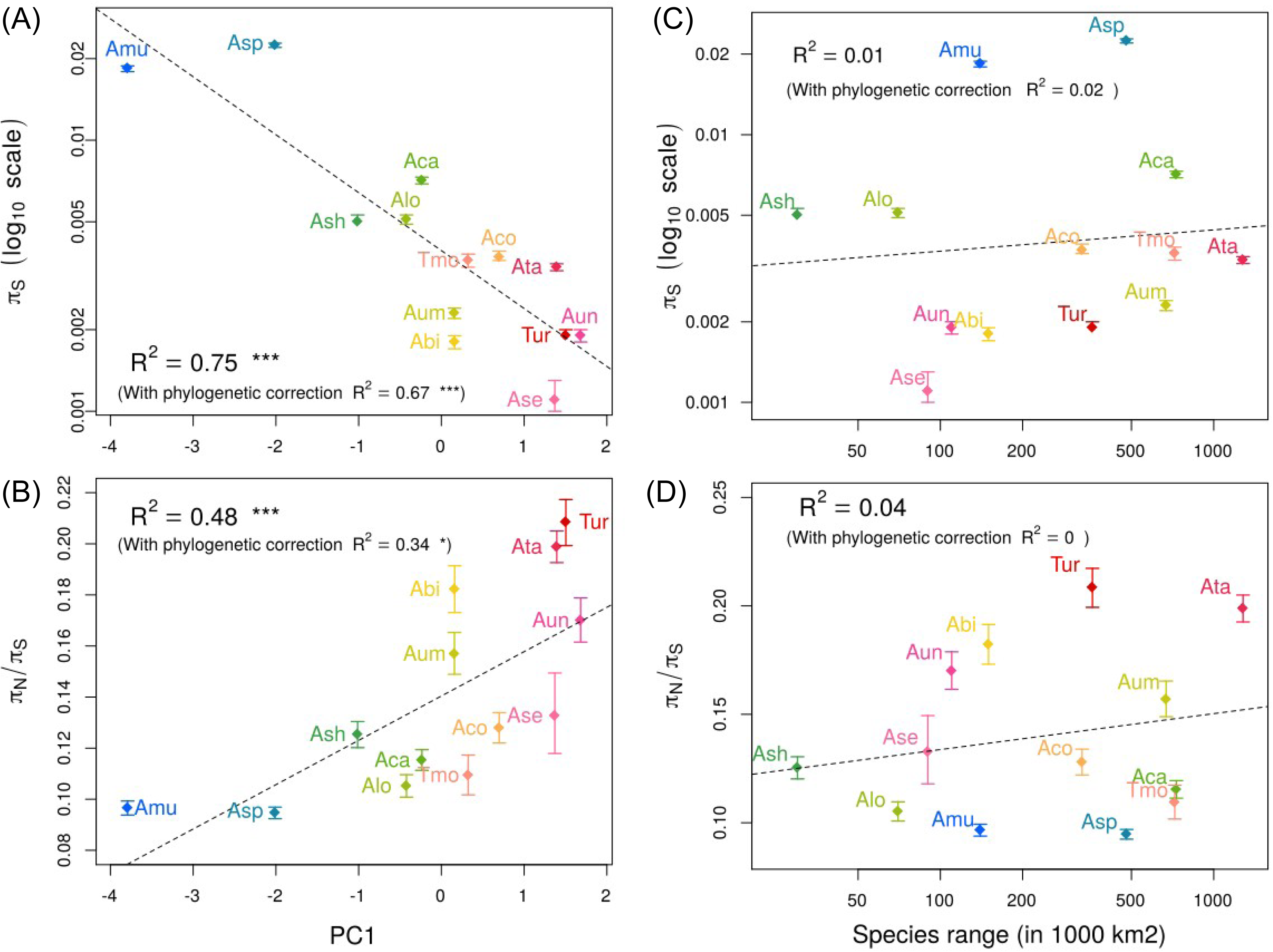
Mating systems traits and global polymorphism patterns. Neutral genetic diversity π_S_ is negatively correlated with the first axis of a PCA of morphofunctional traits associated with reproduction (A), while the efficacy of selection estimated with π_N_/π_S_ ratio is positively correlated with the same PC1 axis (B). Species geographical range estimated from GBIF occurrence data cannot explain the variation observed neither in π_S_ (C) nor π_N_/π_S_ (D).

Selfing is also expected to reduce the efficacy of selection, thus leading to higher accumulation of segregating slightly deleterious mutations in selfing than in outcrossing species (Glémin, 2007). In agreement with this prediction, the efficacy of purifying selection estimated by the ratio of non-synonymous to synonymous polymorphism π_N_/π_S_ (Kimura, 1983) was lower for selfing species (max π_N_/π_S_ value 0.21 for *T. urartu*) than for outcrossing ones (0.09 for *Ae. speltoides*). Similarly to neutral diversity, the efficacy of selection was also significantly explained by the selfing syndrome (R^2^ = 0.48, p-value = 0.0083; with phylogenetic control R^2^ = 0.35, p-value = 0.041; **Fig. 2B**). We further verified that both polymorphism statistics, π_S_ and π_N_/π_S_, also correlated with *F* estimates (**Supplementary Fig. S6**).

We also tested whether species range, used as a proxy of census population size, also correlated with genetic diversity, with widespread species predicted to be more polymorphic than species with restricted geographic distribution. In contrast to the mating system, species range was not correlated with either π_S_ (**Fig. 2C)** or π_N/_π_S_ (**Fig. 2D)**. Such a correlation could be masked by the strong effect of selfing, which is expected to favor species range expansions. For example, *Ae. tauschii* is highly selfing and has by far the largest species range (**Supplementary Fig. S1**). However, a linear model with the two effects showed that the mating system still significantly explained both π_S_ (p-value=0.00004) and π_N_/π_S_ (p-value=0.0157) whereas species range did not (**Supplementary Fig. S7**). Yet, the effect of species range on π_S_ is barely significant (p-value = 0.063), so it is still possible that there is a weak effect that we could not detect with only thirteen species. Overall, these results suggest that the mating system is the main driver of genetic diversity in *Aegilops*/*Triticum* species and overwhelms potential effects of recent population history.

### The effect of linked selection depends on the mating system

We tested the hypothesis that selfing increases the effect of linked selection by comparing polymorphism patterns and recombination along chromosomes. Species-specific recombination maps were not available for most species, so we used the recombination map of the outgroup species *Hordeum vulgare*, which we compared to the recombination maps of the three diploid subgenomes of bread wheat (A, B and D, corresponding to the wild parents *T. urartu*, *Ae. speltoides/mutica*, *Ae. tauschii*). For each focal species, we found that 99% or orthologs with *H. vulgare* mapped on the same chromosome as at least one of the three subgenomes of *T. aestivum*, and 96% to 97% as all three of them. Similarly, 97 to 99% of orthologs with *H. vulgare* belonged to the same recombination category as at least one of the three subgenomes, and 78 to 79% as all three of them (**Supplementary Table S6**).

In what follows, we only show the results with *H. vulgare*, since it provides the further advantage that the outgroup has (on average) the same phylogenetic distance to every *Aegilops*/*Triticum* species, ensuring an unbiased analysis that does not favor species closer to the reference. For comparison, some additional results using the *T. aestivum* subgenomes as reference are given in supplementary material (**Supplementary Table S6** and **Fig. S9**).

In all species, synonymous polymorphism was strongly correlated with recombination and presented a U-shaped pattern along chromosomes more or less mirroring the recombination pattern (see **Fig. 3A** and **Supplementary Fig. S14**). However, the higher the selfing rate, the flatter the relationship (**Fig. 3B**), suggesting a strong effect of the mating system on the relationship between polymorphism and recombination. We verified that the positive relationship between diversity and recombination rate was not merely caused by the mutagenic effect of recombination, by looking at the correlation between synonymous divergence (*D_S_*) with the outgroup (*H. vulgare*) and recombination rate (Kulathinal *et al*., 2008). We found that the magnitude of *D_S_* variation (factor 1.5, **Supplementary Fig. S8**) is much lower than the range of variation observed for polymorphism along the genome (factor 5 to 80, depending on the species, **Fig. 3B**). If a mutagenic effect of recombination cannot be ruled out, it is clearly insufficient to explain the magnitude of the correlation between π*_S_* and recombination. This mere observation suggested that linked selection could strongly reduce π*_S_* by at least one or two orders of magnitude.

**Figure 3.**
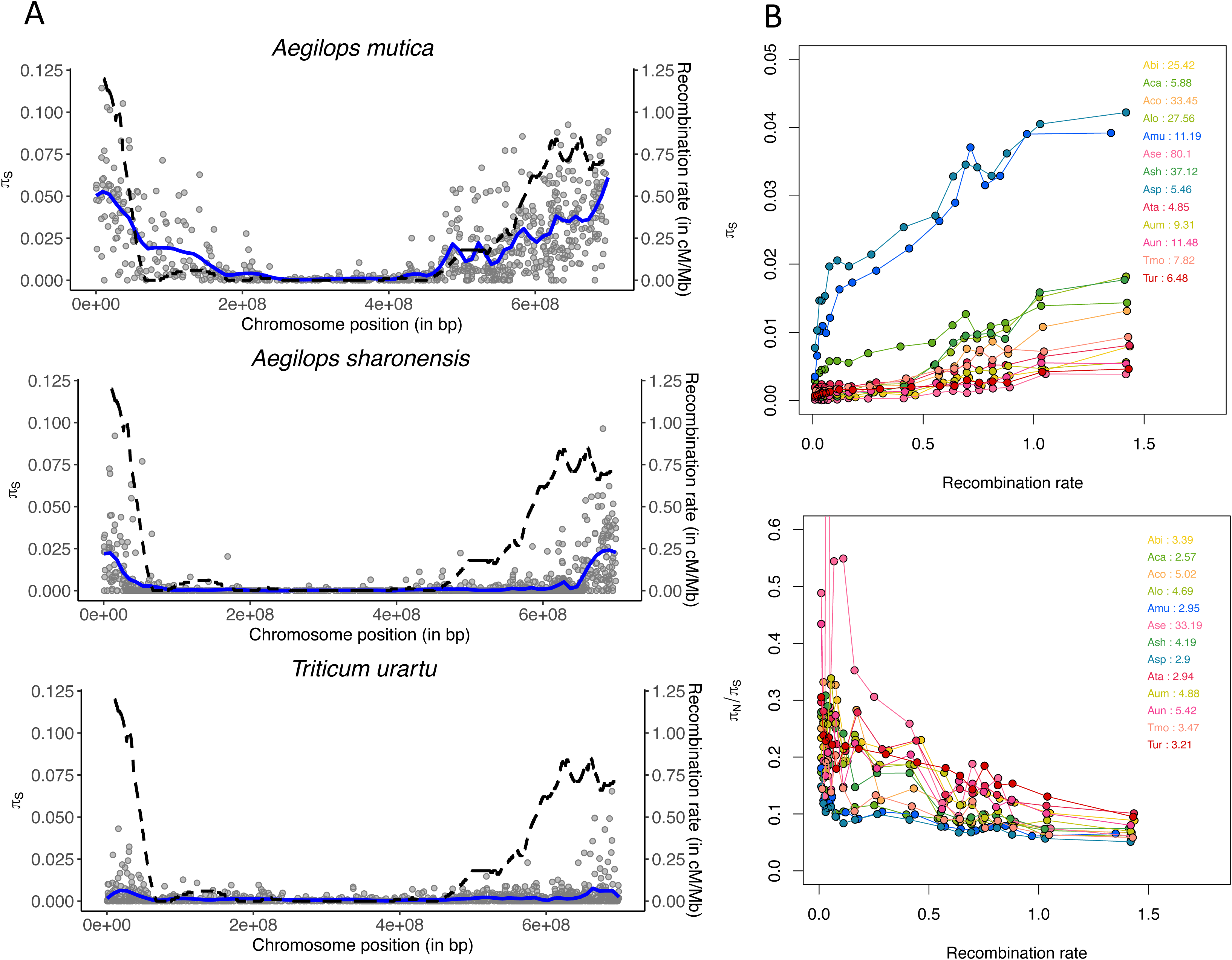
Recombination and patterns of genetic polymorphism across the genome. (A) Mean synonymous diversity (π_S_) along chromosome 3 for three species with contrasted mating system: self-incompatible (*Ae. mutica*), mixed-mating (*Ae. sharonensis*) and highly selfing (*T. urartu*); same scales for the three species. Each point corresponds to a contig mapped on the *Hordeum vulgare* genome. The blue line is the loess fitting function (degree=2, span=0.1). The dashed black line indicates *H. vulgare* recombination map (in cM/Mb). (B) π_S_ and π_N_/π_S_ as a function of recombination rate. Contigs have been grouped in 20 quantiles of recombination. The value associated with each species corresponds to the ratio between the highest and the lowest value among the quantiles: from about 5 to 80 for π_S_ and from 2.5 to 33 for π_N_/π_S_. Curves correspond to loess fitting functions (degree = 2, span = 0.2).

To quantify more directly the effect of linked selection, we fitted a model similar to Corbett-Detig *et al*. (2015) and Elyashiv *et al*. (2016) that we adapted to partially selfing species but only considering background selection (see **Supplementary Table S6** for full results). From the fit of the model, we obtained the maximum π*_S_* that could be reached in the absence of linked selection, π_max_, which ranged between 0.028 to 5.82 (**Fig. 4**). Note that, here π_max_ = 4*N*_e_max_*u*, so can be higher than 1. We estimated that linked selection reduced π_S_ by 3.5 in *Ae. speltoides* and 5.6 in *Ae. mutica*, the two self-incompatible species. For other species, π_S_ was reduced by a few tens or even a few hundreds (from 7 to 888), but without a clear relationship with the mating system. In contrast to π_S_, π_max_ did not correlate with PC1. Surprisingly, it correlated negatively with species range, but the correlation was mainly driven by the species of the *Sitopsis* section and was no longer significant after phylogenetic correction (**Fig. 4**). It can be difficult to properly fit a realistic linked selection model for selfing species and the results can be sensitive to the fact that we did not use the reference genome of each species. In particular, some π_max_ values were very high and could be overestimated, but fitting the model using the three subgenomes A, B and D of *T. aestivum* gave similar results (**Supplementary Fig. S9**). Overall, although they must be viewed with caution, the results strongly suggested that linked selection is a main driver of the effect of selfing on genetic diversity whereas species range has only a minor effect, and if any, not in the predicted direction. When recombination maps will be available in all species, it will be possible to re-assess this result.

**Figure 4.**
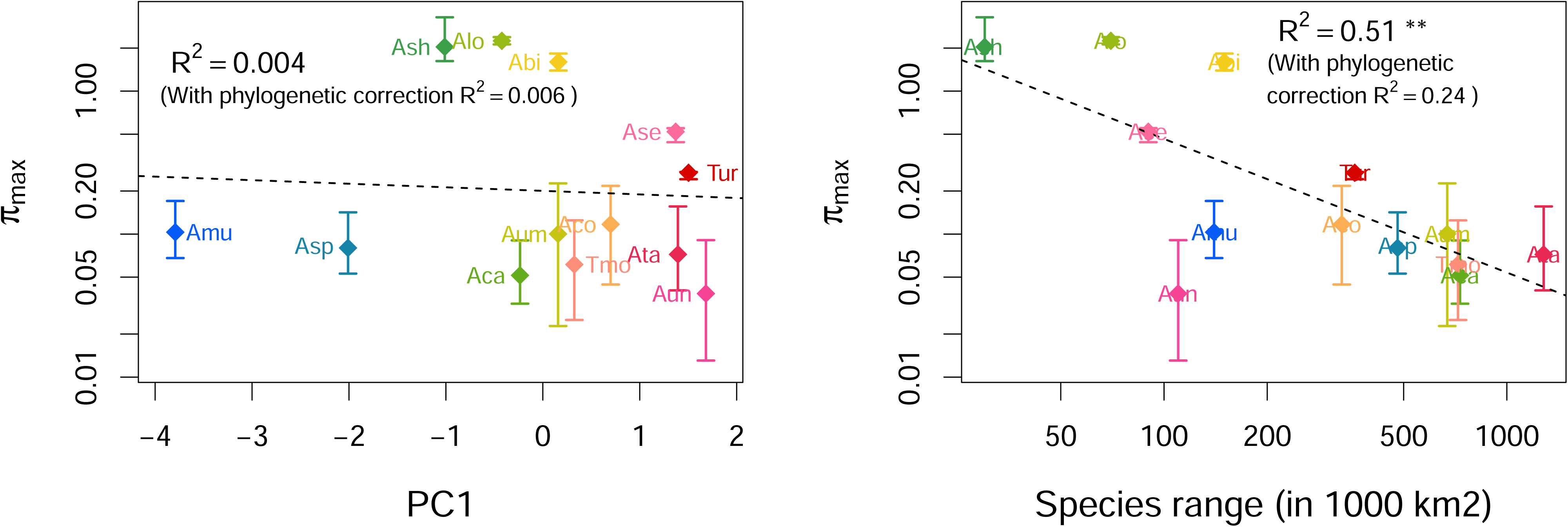
Relation of estimated maximum polymorphism in absence of linked-selection and mating system. π_max_ does not correlate with the first axis of the PCA of morphofunctional traits (left). π_max_ correlates negatively with species range (right), but the correlation is mainly driven by the species of the *Sitopsis* section and is no longer significant after phylogenetic correction.

### Deleterious mutations accumulate and adaptation is reduced under selfing and low recombination

Another central prediction of the effect of genetic linkage is that selection should be less efficient in genomic regions of low recombination, which can extend genome wide in highly selfing species. In agreement with this expectation, the efficacy of purifying selection at the genome wide level clearly decreased with the selfing rate (**Fig. 2B**). All species also showed a negative relationship between recombination rates and the π_N_/π_S_ ratio (**Fig. 3B**), indicating that purifying selection was more efficient in highly recombining regions. More precisely, the π_N_/π_S_ ratio sharply dropped with increasing recombination in outcrossing species but more and more smoothly with increasing selfing rate, which supports the prediction that reduced selection efficacy extended to larger genomic regions in selfing species.

The π_N_/π_S_ ratio is a rather crude proxy for the efficacy of purifying selection, and can be affected by several factors such as non-equilibrium population dynamics that can lead to spurious signature of relaxed selection (Brandvain and Wright 2016). To better characterize how selection efficacy varies with mating system and recombination, we estimated the full distribution of fitness effects (DFE) of mutations – i.e. including both deleterious and beneficial mutations – using the *polyDFE* method (Tataru *et al*., 2017). This approach leverages information from unfolded synonymous and non-synonymous site frequency spectra to infer the DFE of each species. It takes into account factors that can distort the SFS such as non-equilibrium demography and linked selection, in addition to potential polarization errors. The method requires a sufficient number of chromosomes sampled (say >10), so we applied it only to the four species with the largest sample sizes, which correspond to the extreme mating systems used to calibrate the selfing syndrome: the two self-incompatible *Ae. speltoïdes* and *Ae. mutica* and two highly selfing *Ae. tauschii* and *T. urartu*. In agreement with the π_N_/π_S_ ratio analysis, we found that the two selfers suffered from a higher load than the two outcrossers, with 22% to 25% of mutations not being efficiently selected against (–10 < *Ne s* < 0), versus only 9 to 15% in the outcrossers (**Fig. 5**). We also found a strong difference between regions of low and high recombination (higher vs lower than the median) for all species. However, the difference was stronger in outcrossers (more than two-fold) than in selfers (30-60% difference only) (**Fig. 5**). Interestingly, purifying selection appeared as efficient in low-recombination regions of outcrossing genomes than in high-recombination regions of highly selfing genomes (**Fig. 5**).

**Figure 5.**
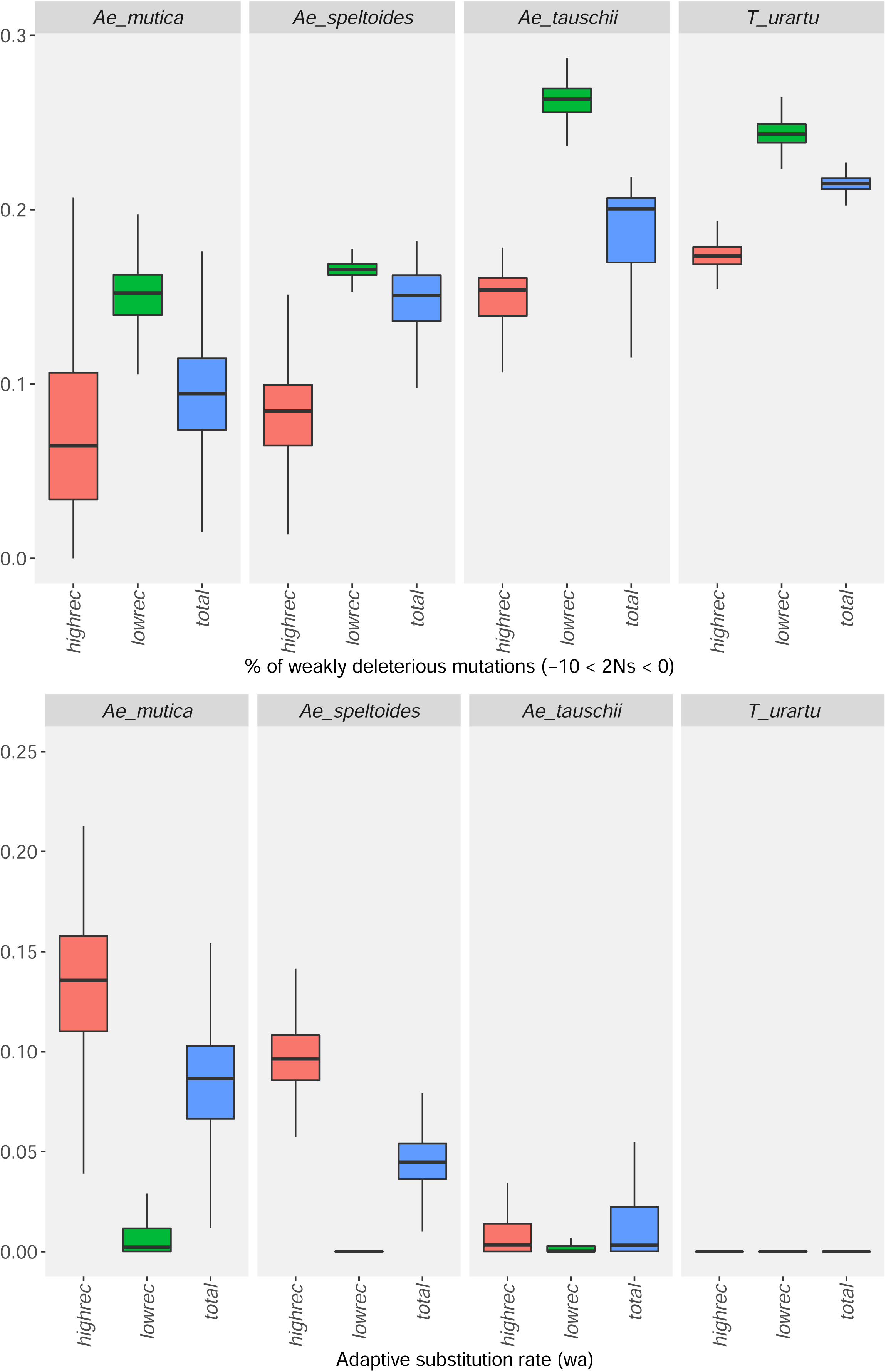
Mating systems, recombination and the efficacy of selection. The distribution of fitness effect of mutations was estimated in two self-incompatible species (*Ae. mutica* and *Ae. speltoides*) and in two highly selfing species (*A. tauschii* and *T. urartu*) at the whole genome scale (blue) and for regions of low (green) and high (red) recombination. (A) Proportion of weakly selected deleterious mutations (–10 < 2*N*_e_*s* < 0). (B) Rate of adaptive substitution, i.e. the expected *D*_N_/*D*_S_ ratio due to the flux of beneficial mutations.

Another striking result is that we estimated an adaptive substitution rate not different from zero in the two selfers, but a quite high value in the two outcrossers and only in highly recombining regions (**Fig. 5**). We verified that this signature of positive selection was not due to the spurious effect of GC-biased gene conversion (**Supplementary Fig. S10**), which is known to happen in recombining regions of grass genomes (Muyle *et al*. 2011). Overall, selection appeared to be much less efficient, both on beneficial and against deleterious mutations, in low-recombination regions of outcrossing species, and throughout the genome of highly selfing ones. When the proportion of weakly deleterious mutations is estimated with the DFE-alpha method (Keightley and Eyre-Walker 2007) it was shown that selfing could overestimate it (Gilbert et al. 2022). Here, we used polyDFE, which was claimed to be less sensitive to linked selection effects (Tataru et al. 2017). As a control, we run simulations with linked selection and varying degree of selfing and applied polyDFE. In contrast with DFE-alpha (see Gilbert et al. 2022), we found that polyDFE tended to underestimate the proportion of weakly deleterious mutations (**Supplementary Fig. S11).** We also found that the method did not estimate spurious signatures of beneficial mutations in outcrossers. Overall, the results were conservative to the effects of linked selection and selfing.

## Discussion

We compared the patterns of genetic diversity and selection across the genome in thirteen species with contrasted mating systems. We found far less polymorphism and far less selection efficacy in selfing than in outcrossing species, as observed in previous studies (e.g., Glémin *et al*., 2006; Hazzouri *et al*., 2013; Slotte *et al*., 2013; Barrett *et al*., 2014; Burgarella *et al*., 2015; Chen *et al*., 2017; Laenen *et al*., 2018). We also showed that these genomic effects depend on the interplay between linked selection and mating systems, and vary with self-fertilization rates. For this, we leveraged a study design tailored to go beyond global patterns and decipher their underlying causes. First, differently from previous general comparisons among plant species (Glémin *et al*., 2006; Chen *et al*., 2017), we compared related species with similar life history traits, ecology and genomic features, which allows more direct testing of the effect of the mating system. Second, we investigated all species of a clade covering a large range of mating systems, including intermediate mixed mating species, whereas previous studies addressing sister (or closely related) species mainly focused on extreme outcrossing vs selfing comparisons (e.g., Hazzouri *et al*., 2013; Slotte *et al*., 2013; Burgarella *et al*., 2015; Teterina *et al*., 2023). Third, by using recombination maps we quantified the effect of linked selection in a comparative way.

Linked selection appears to be a main mechanism shaping levels of diversity in wild wheats, as it can reduce polymorphism to three to five-fold in outcrossing species (**Fig. 3B**) and to one or two orders of magnitude in selfing species. This is in agreement with but higher than observed by Corbett-Detig *et al*. (2015), who found a quantitatively limited effect of linked selection except in selfing species (see the re-analysis of Coop 2016). These quantitative values must be viewed with caution because it is difficult to properly fit a model of linked selection in selfing species. In addition to the complexity of the interaction between recombination, selfing and selection, it is not clear whether the species scale or a more local population scale is the most relevant. Another expected consequence of linkage and selfing is a reduction in selection efficacy, both against deleterious and in favor of beneficial mutations. We observed a striking contrast between regions of high and low recombination in self-incompatible species, with twice more weakly selected deleterious mutations and no beneficial mutation expected to fix in regions of low recombination (**Fig. 5, Supplementary Fig. S10**). In highly selfing species, instead, recombination had a weaker effect and only for deleterious mutations (**Fig. 5)**. Interestingly, the regions of high recombination in selfers exhibited a similar amount of weakly deleterious mutations as regions of low recombination in outcrossers, and we detected no signature of adaptive evolution at all in the two highly selfing species. Simulations showed that these results were not artefactual and may even underestimate the effect of selfing (**Supplementary Fig. S11**). Overall, the cross comparison between mating systems and recombination levels clearly showed that the main quantitative effect of selfing is due to high linkage and linked selection.

Although our studied species share life history traits and have similar ecology, other factors than the mating system could affect genetic diversity, for example, factors generating contrasted geographic ranges unrelated to mating systems. However, we showed that the geographic range has no effect on polymorphism patterns (**Fig. 2, Supplementary Fig. S7**). More generally, we cannot exclude that other factors could play a role, but they could hardly generate the very strong relationship between the mating system and genetic diversity we observed. Strikingly, this strong relationship holds despite the use of an indirect measure of the selfing rates through phenotypic proxies.

Our results help better understand the evolution of selfing species. In the short term, selfing is known to recurrently evolve from outcrossing, depending on reproductive assurance and gene transmission advantage balanced by inbreeding depression, which can be partly purged during such transitions. In the long run, selfing lineages tend to diversify less than outcrossing ones and selfing is considered an evolutionary dead-end, likely because of higher probability of extinction (Stebbins, 1957; Igic *et al*., 2008; Goldberg and Igić, 2012). The very causes of higher extinction rates in selfers remain unclear, but increased load and loss of genetic diversity and adaptive potential are possible drivers. However, the pace at which the effects of selfing manifest is still poorly known, although it is likely rapid as in the selfing *Capsella rubella* recently derived from the self-incompatible *C. grandiflora* (Slotte *et al*., 2013) or within the species *Arabis alpina* among populations with contrasted mating systems (Laenen *et al*., 2018).

In the *Aegilops*/*Triticum* genus, several instances of evolution towards different degrees of selfing likely occurred in a short evolutionary time period, which manifested by wide variations in floral traits associated with specific genetic diversity patterns. At the short phylogenetically scale we studied, we thus found a clear signature of the joint evolution of morphological traits and population genomic patterns, suggesting that the negative effects of selfing manifests rapidly. It is tempting to propose that the strong and rapid deleterious effects of selfing we detected will accelerate the extinction of the most selfing lineages. However, so far there is no approach to properly test this hypothesis (Wright *et al*., 2013; Glémin *et al*., 2019a). Genomic degradation certainly accompanies the transition towards selfing, but we still do not know whether it is the ultimate cause of selfing lineages extinction.

Finally, our results also bear more general implications about the central question of the determinants of genetic diversity, beyond the case of selfing species. In line with previous comparative analyses of polymorphism at the genome scale (Romiguier *et al*., 2014; Chen *et al*., 2017), we showed that life history traits, here the mating system, are a much stronger determinant of genetic diversity than proxies for census population sizes, here species range. Species range had no effect either after globally controlling for the mating system (**Supplementary Fig. S7),** or after removing specifically the effect of linked selection (no significant correlation between species range and π_max_ or, if so, in the unpredicted direction). In contrast to previous studies, however, the range of genetic diversity is particularly large despite species being closely related and recently diverged (about 6 MYA), with similar genomes and life-history traits (wind-pollinated annual herbs) except mating system. In these wild wheats, species nucleotide polymorphism varies with a factor 20 (from 0.0011 to 0.022). For comparison, a clade of selfing and outcrossing *Caenorhabditis* nematodes species diverged less than 30 MYA show even wider disparities in polymorphism, up to a factor 80 (Cutter 2008; Li *et al*. 2014). In contrast, in a butterfly family that diverged around 120 MYA, with four-fold variation in body mass and two-fold variation in chromosome numbers, only a factor ten was observed (from 0.0044 to 0.043) (Mackintosh *et al*., 2019). Similarly, among 28 species widely covering the seed-plant phylogeny and life forms (from annual herbs to trees), the observed range was only slightly higher than in wild wheats, with a factor 28 (from 0.00064 to 0.018) (Chen *et al*., 2017). This points to mating systems as a main determinant of variation in genetic diversity among species.

However, variation in genetic diversity is still narrower than predicted from variation in species range, which varies by a factor 500 here. This is in line with the “Lewontin’s paradox”, the general observation that genetic diversity varies much less across species than census size does (Lewontin, 1974; Charlesworth and Jensen, 2022). Variation in π_max_ is around 60, so higher than variation in π_S_, around 20, but only by a factor three. Despite its strong effect, linked selection is thus unlikely to explain alone the limited range of variation in π_S_, in agreement with previous results (Corbett-Detig *et al*., 2015; Coop, 2016; Buffalo, 2021; Charlesworth and Jensen, 2022).

## Acknowledgements

We thank G. Sarah, Y. Holtz and P. Joncour for help with bioinformatic analyses and T. Bataillon, M. Lascoux and D. Schoen for helpful discussions and suggestions on the manuscript. The work was funded by the French Agence Nationale de la Recherche (ANR) (ANR-11-BSV7-013-03). CB has also received funding from the European Union’s Horizon 2020 research and innovation programme under the Marie Skłodowska-Curie grant agreement No. 839643. The authors declare no conflicts of interest.

## Data and code availability

Custom R and bash codes used for the analyses are available on https://github.com/sylvainglemin/ms-rec-triticeae along with input files. Software for genotype calling (reads2snps v. 2.0.64, ORF_extractor.pl) and polymorphism estimates (dNdSpNpS v.3) are available at https://kimura.univ-montp2.fr/PopPhyl/index.php?section=tools. Morphological traits measures are provided as Supplementary Tables S2 and S3.

Filtered and cleaned sequence alignments to perform polymorphism analyses are available at https://bioweb.supagro.inra.fr/WheatRelativeHistory/index.php?menu=downloadMating. Raw data are deposited at the Sequence Read Archives (SRA) under project PRJNA945064 (submission number SUB12943046).

## Supplementary Information

Figures S1 to S14

Supplementary tables:

Supplementary Table S1. Passport information for the accessions of Triticum/Aegilops species and outgroups used in this study.

Supplementary Table S2. Dataset 1: Morphofunctional traits linked to reproduction measured in 13 Triticum/Aegilops species and three outgroups.

Supplementary Table S3. Dataset 2: Morphofunctional traits linked to reproduction measured in 13 Triticum/Aegilops species and three outgroups where missing data were imputed with package missMDA under R environment (see Materials and Methods).

Supplementary Table S4. List of morphological traits measured and used for the analysis.

Supplementary Table S5. Sequencing and genome summary statistics per species (13 Triticum/Aegilops species and three outgroups).

Supplementary Table S6. Comparison of genes mapped on the Hordeum genome with the characteristics of the three Triticum sub-genomes (A, B and D).

Supplementary Table S7. Results of the linked-selection model.

