## Supplemental Figures for "Mating systems and recombination landscape strongly shape genetic diversity and selection in wheat relatives"

*Manuscript:*

**This PDF file includes Supplementary Figures S1 – S14**

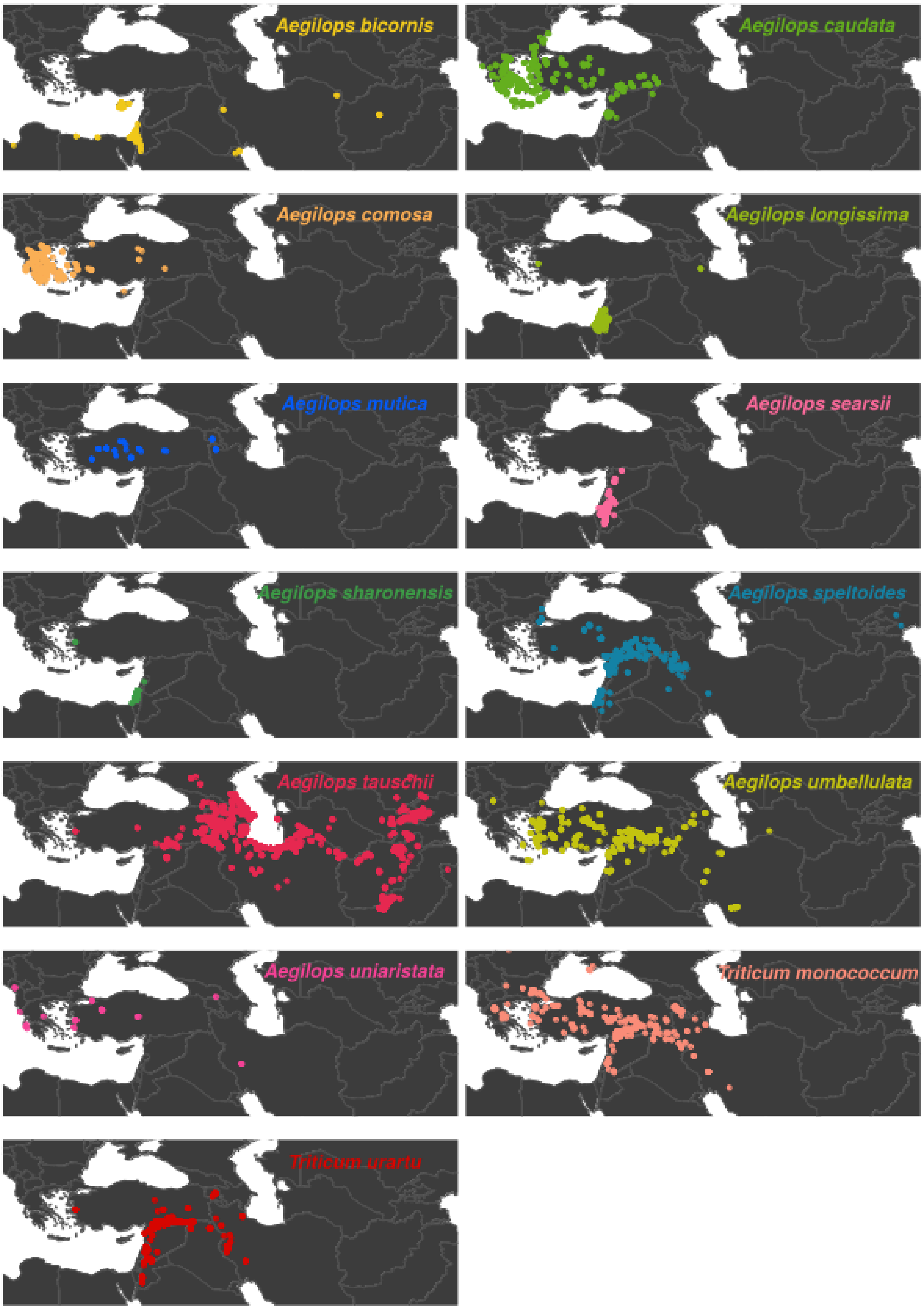


**Figure S1.** Geographic distribution of the 13 diploid *Aegilops*/*Triticum* species. Each dot corresponds to an observation retrieved from GBIF (http://www.gbif.org). Color code corresponds to Figure 2 of the main text.

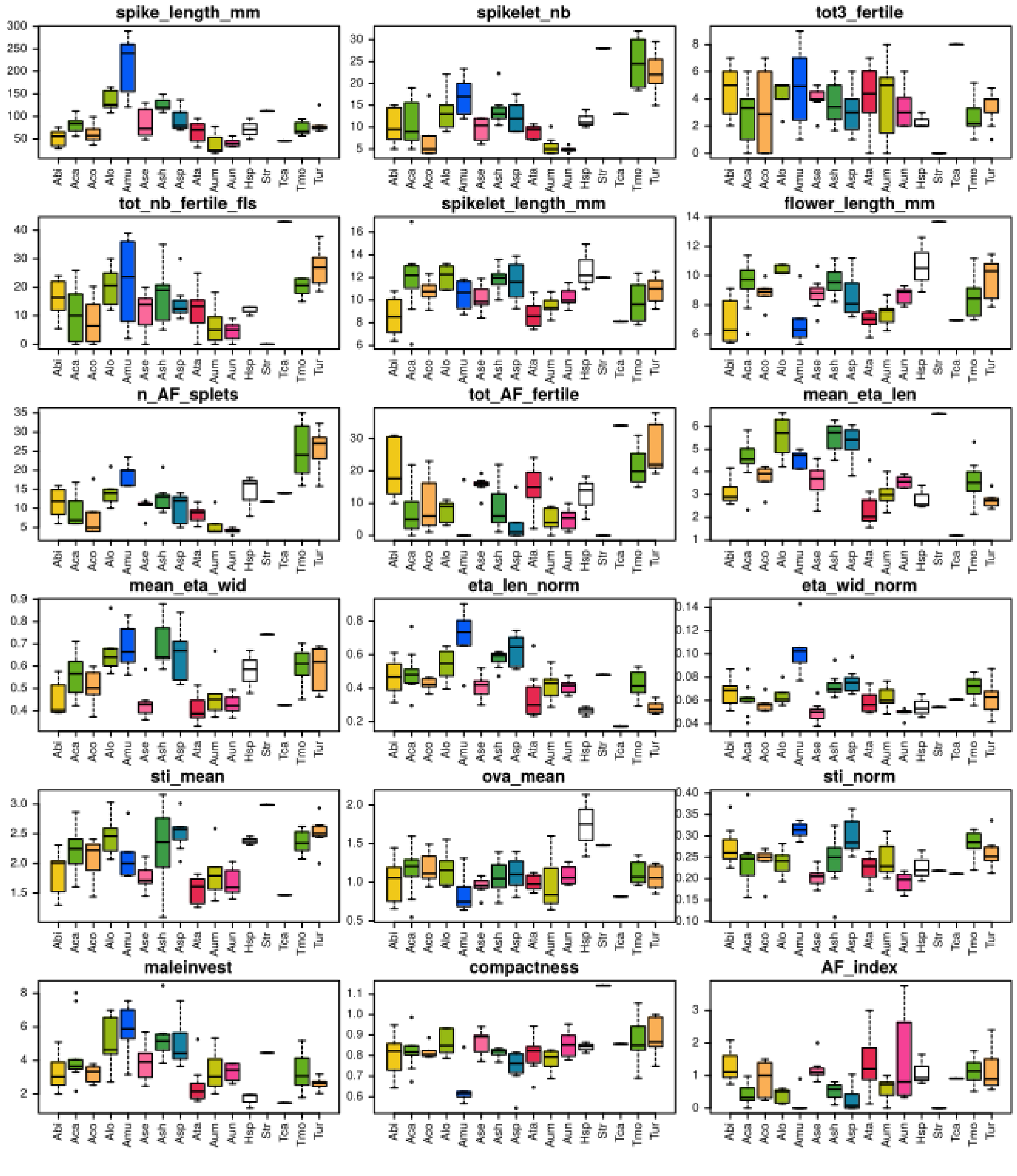


**Figure S2.** Distribution of 18 morphological traits linked to reproduction in the 13 *Aegilops*/*Triticum* species. Missing data were imputed with a PCA based approach implemented in missDA under R environment (see Material and Methods and Supplementary Tables S3 for the meaning of species code and trait names). Colors correspond to Figure 2 of the main text.

(A)

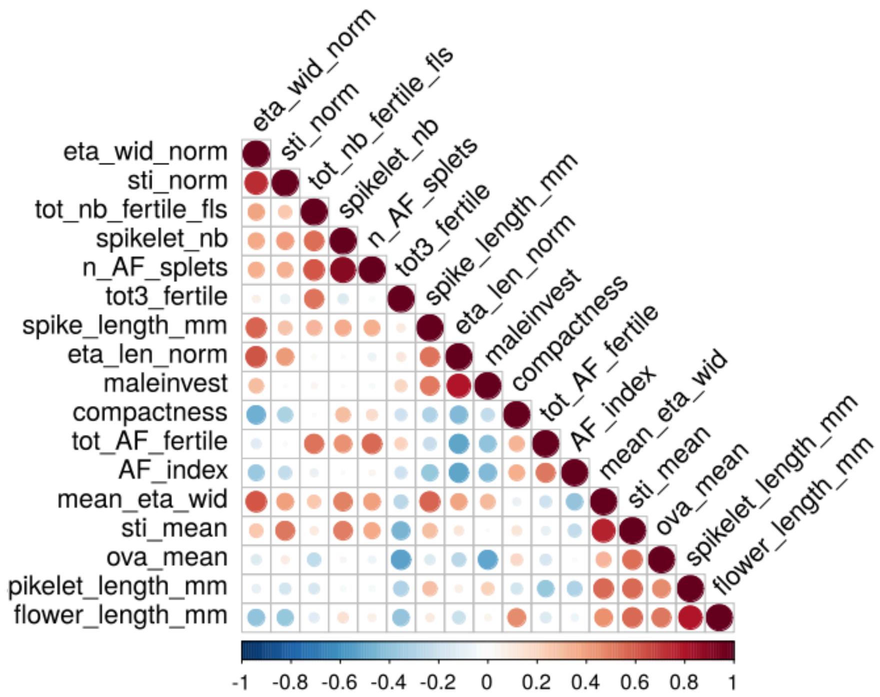


(B)

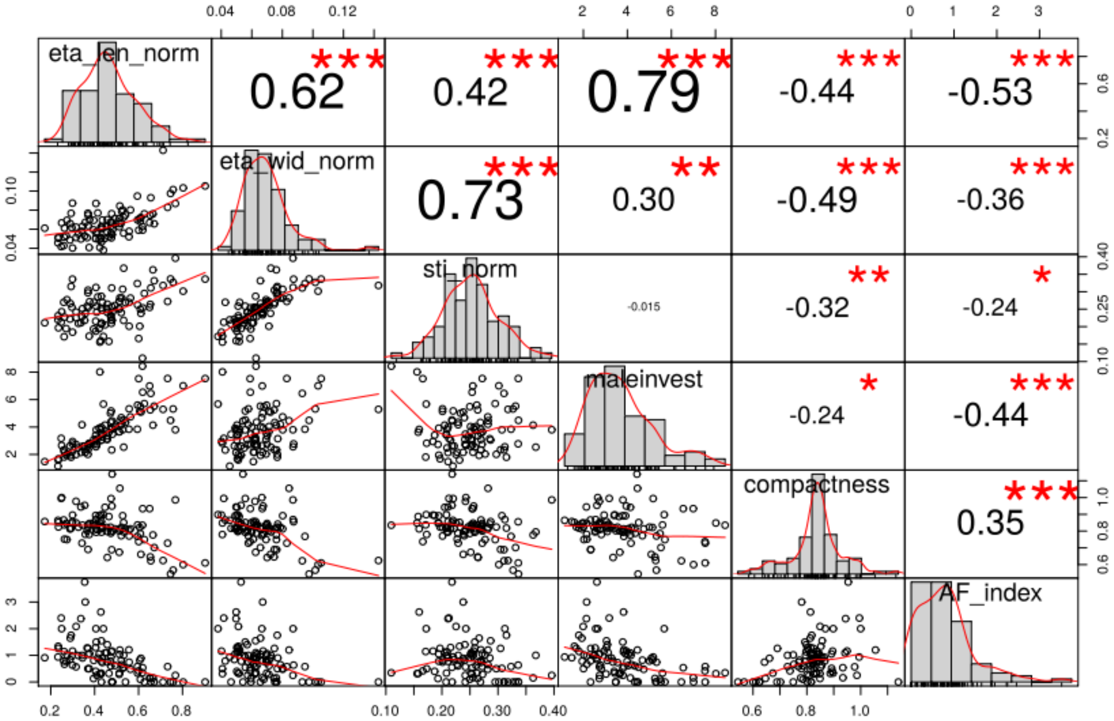


**Figure S3.** Pairwise correlations among 18 morphological traits related to reproduction and measured in the 13 diploid *Aegilops*/*Triticum* species: (A) all traits, (B) traits used in the principal component analysis. Missing data were imputed with a PCA based approach implemented in missDA under R environment (see Material and Methods and Supplementary Table S3 for the meaning of trait names).

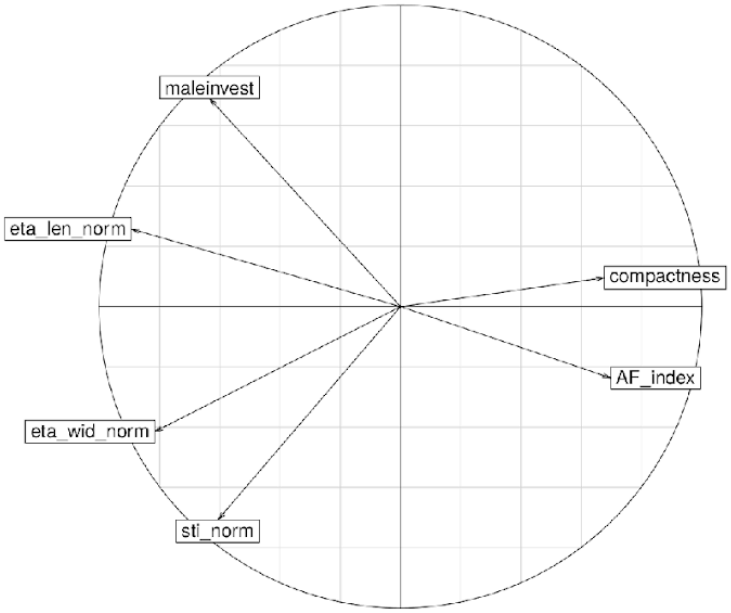


**Figure S4.** Contribution of six selected morphofunctional traits to a Principal Component Analysis describing the morphological selfing syndrome in the 13 diploid *Aegilops*/*Triticum* species. Missing data were imputed with a PCA based approach implemented in missDA under R environment (see Material and Methods and Supplementary Table S3 for the meaning of trait names).

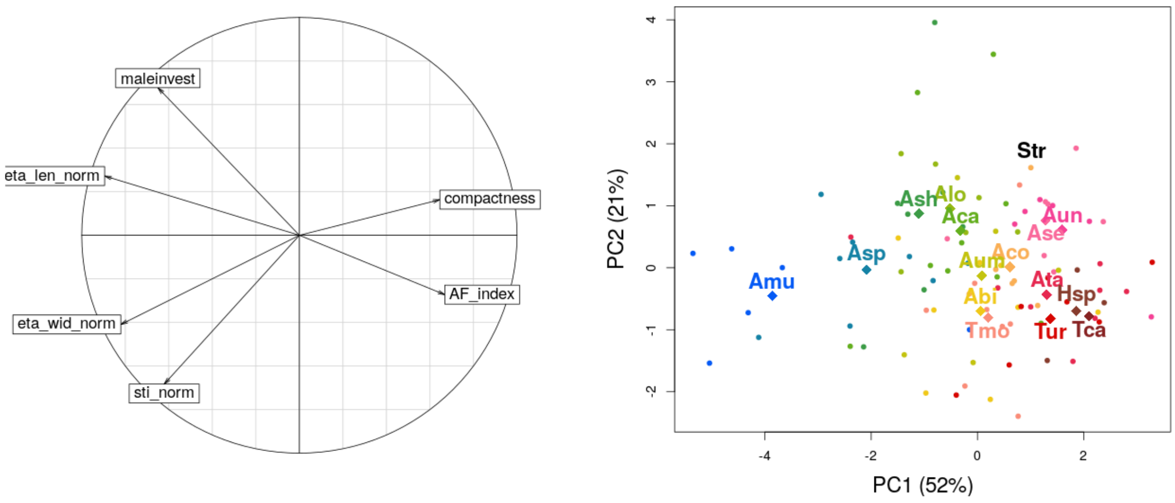


**Figure S5.** Results of PCA of morphological traits of reproductive organs when all species (including outgroups) are used for missing data imputation and PCA. Including the outgroups does not change the relative position of *Aegilops*/*Triticum* species within the PC1. Species code is the same than Supplementary Table S3.

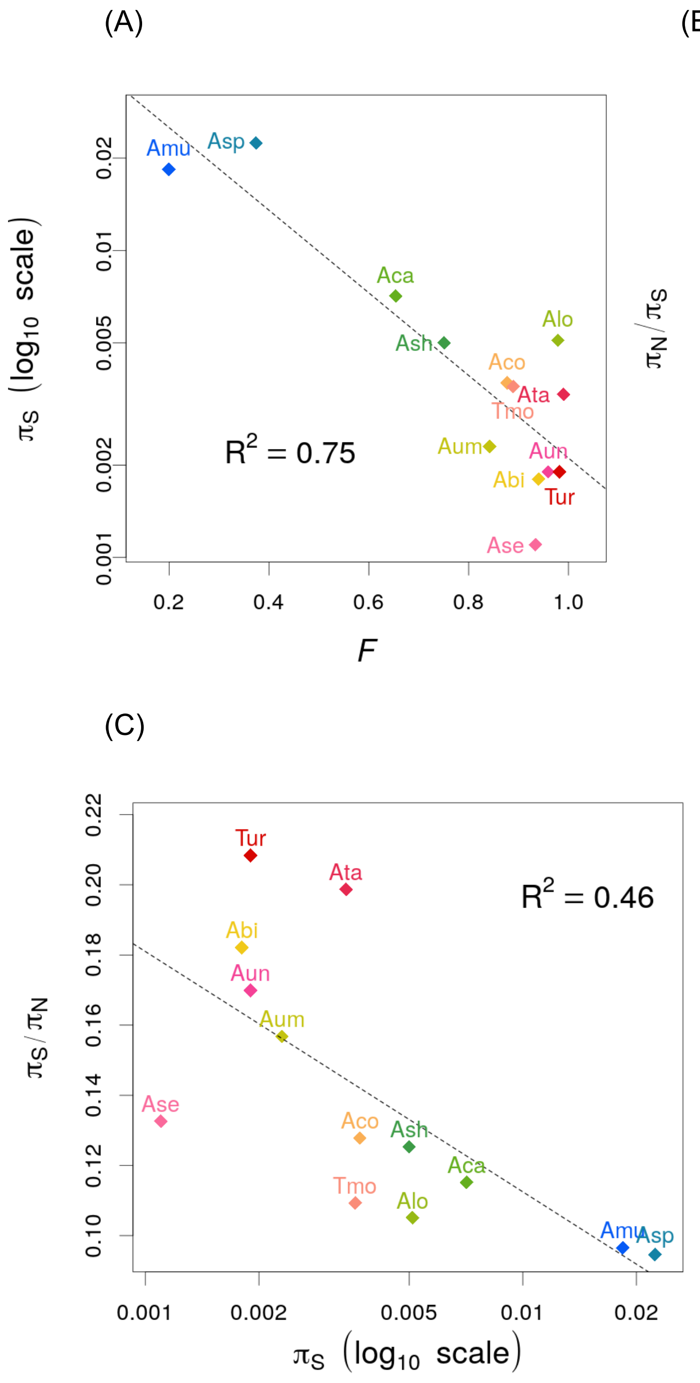


**Figure S6.** Correlation of polymorphism estimates $\pi_S$ (A) and $\pi_N/\pi_S$ (B) with the inbreeding coefficient *F*, and of $\pi_N/\pi_S$ with $\pi_S$ (C).

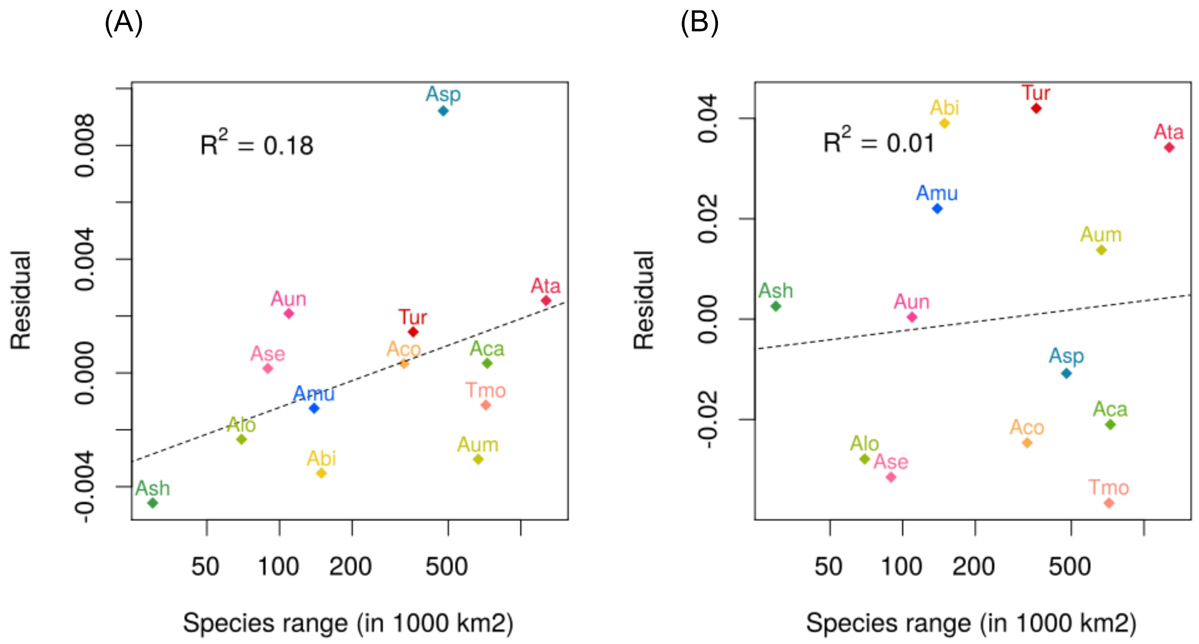


**Figure S7.** Joint effect of mating system and species range on $\pi_S$ (A) and $\pi_N/\pi_S$ (B). Multiple regressions have been performed: log($\pi_S$) ~ mating_system + species range (mating_system: p-value < 0.0001, species_range: p-value = 0.063). $\pi_N/\pi_S$ ~ mating_system + species range (mating_system: p-value = 0.016, species_range: p-value = 0.67). The plots represent the residuals, after removing the effect of mating system, as a function of species range.

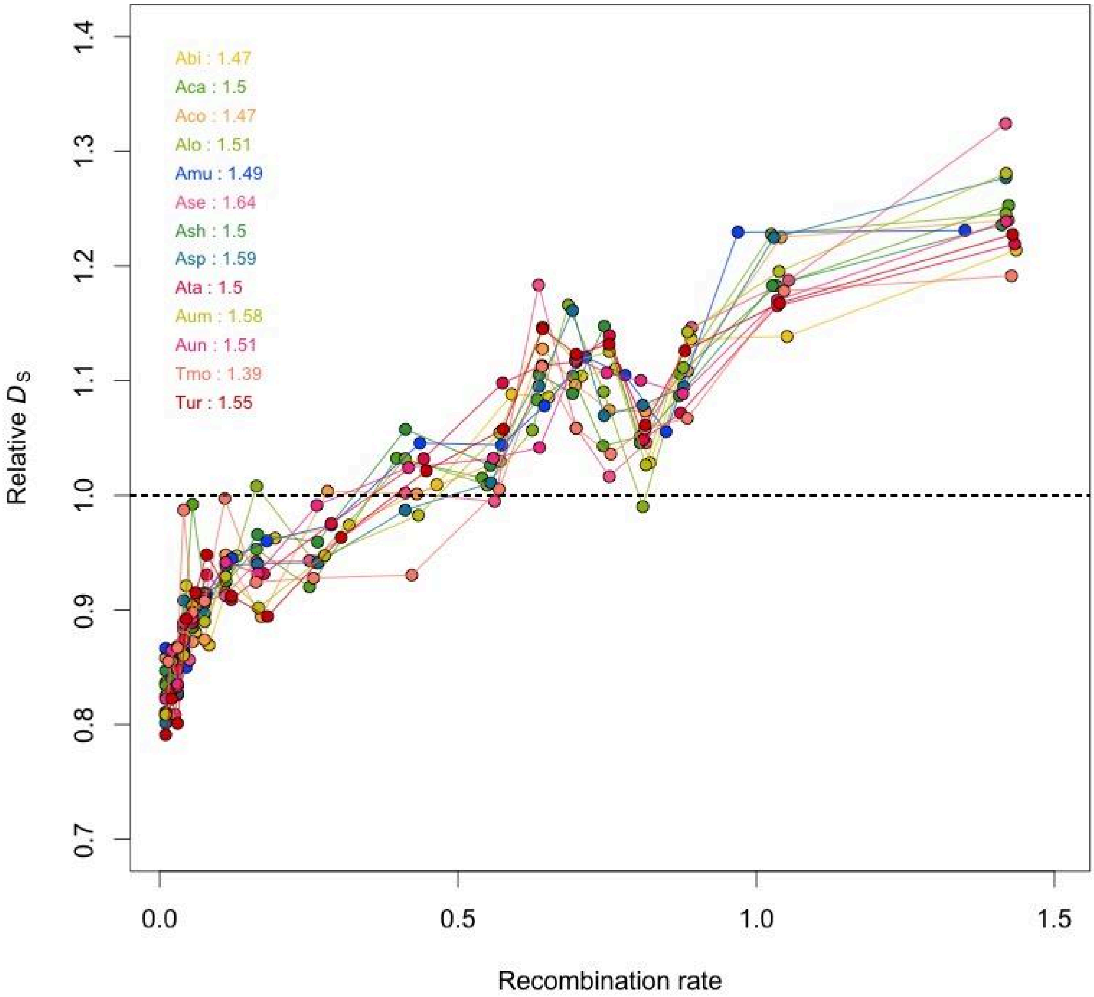


**Figure S8**: Relative $D_S$ ($D_S$ divided by the mean genome wide $D_S$) as a function of recombination rate. Contigs have been grouped in 20 quantiles of recombination. Each color corresponds to a species (same code as in the main text). The value associated with each species corresponds to the ratio between the highest and the lowest value among the quantiles. Curves correspond to loess fitting functions (degree = 2, span = 0.2). Variation in $D_S$ (factor 1.5) is much lower than the range of variation in $\pi_S$ (factor 5 to 80, depending on the species, see main text Figure 3).

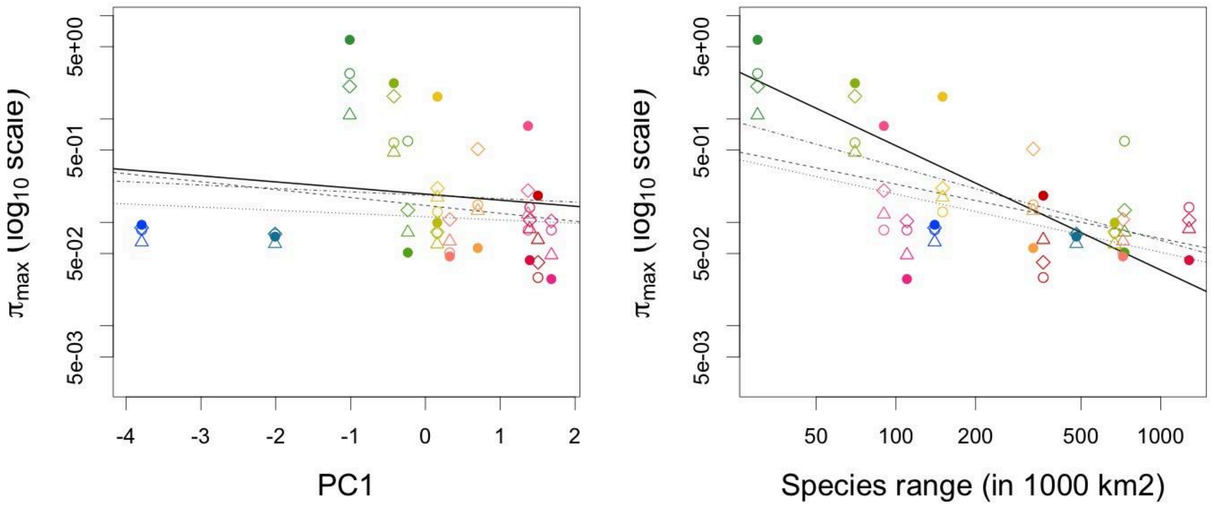


**Figure S9.** Fitted $\pi_{max}$ as a function of PC1 and species range, using either the *Hordeum* genome as a reference (as in the main text): full circles and plain regression line, or with the three subgenomes of *Triticum aestivum*: A genome: open circle and dashed line, B genome: open triangle and dotted line, D genome: open diamond and dotted-dashed line. Confidence intervals are not presented not to overweight the figure.

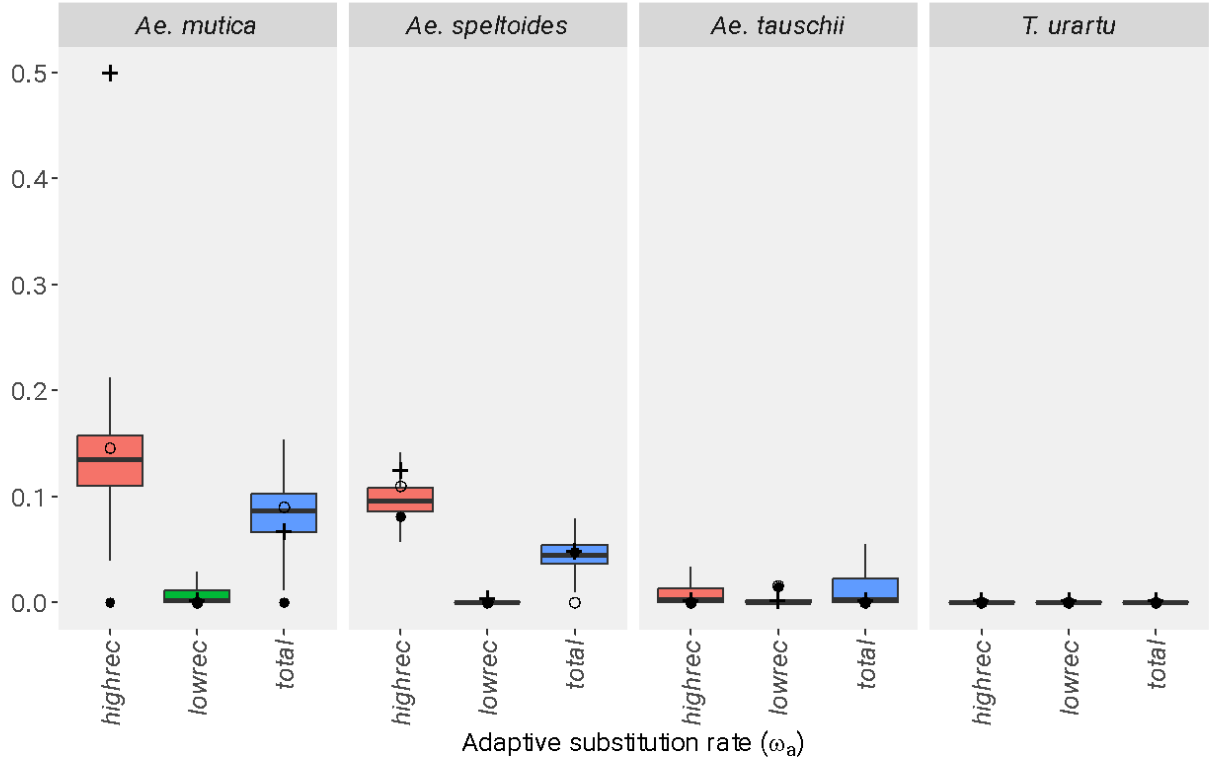


**Figure S10.** Adaptive substitution rate estimated with *polyDFE* for different classes of AT/GC mutations (point estimates): GC-conservative (+), AT➔GC (o) and GC➔AT (●). gBGC may perturb the estimation of DFE parameters but the signature of positive selection in recombining regions of outcrossing species is not due AT➔GC mutations. In particular, the strongest signatures are observed for GC-conservative mutations.

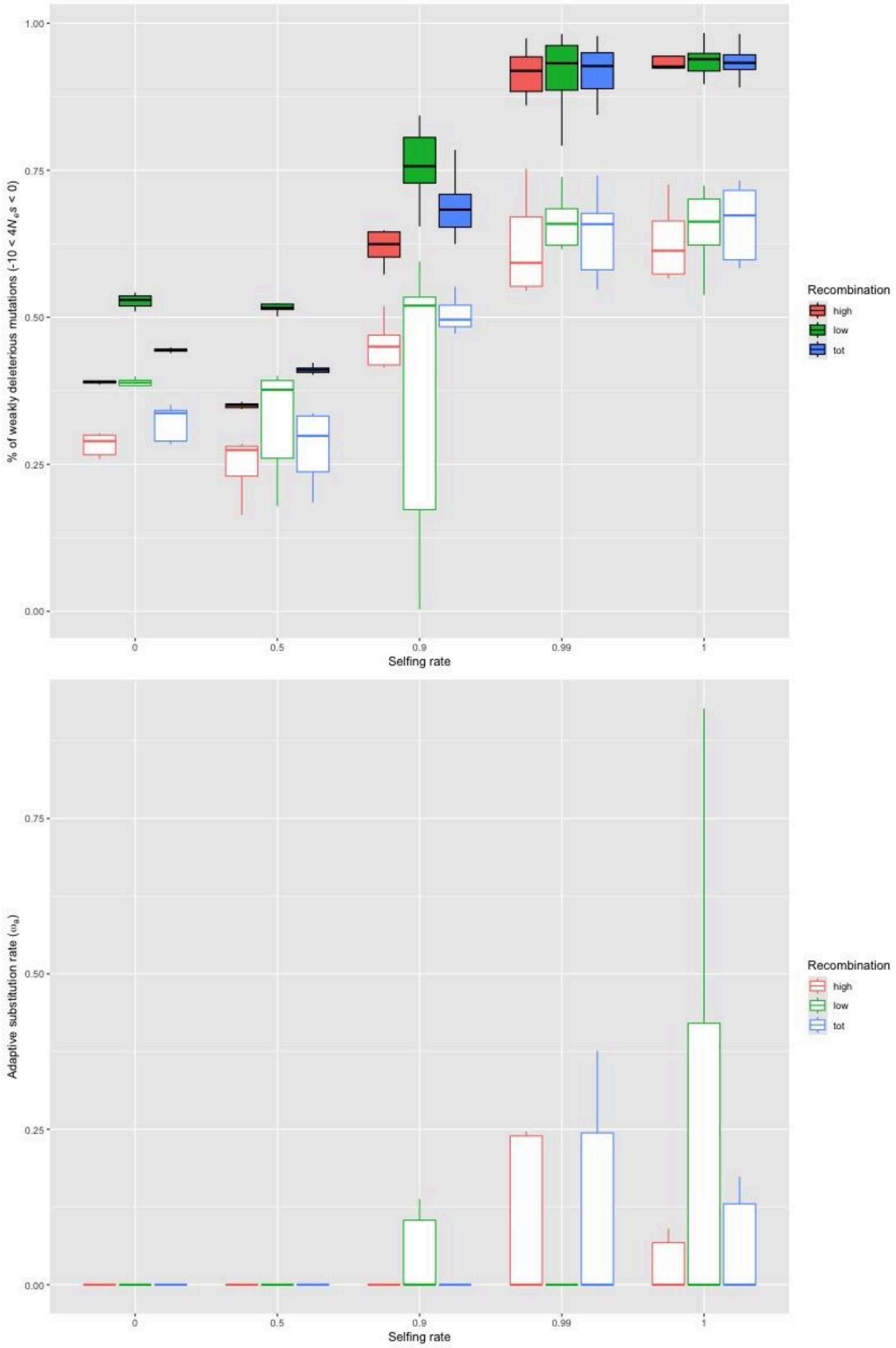


**Figure S11** Estimation of the proportions of weakly deleterious (top) and adaptive (bottom) mutations in simulated datasets with background selection and varying degree of selfing. The filled boxplots correspond to expectations and the open ones to estimation with polyDFE. For the adaptive substitution rate, the expectation is zero as only deleterious mutations have been simulated.

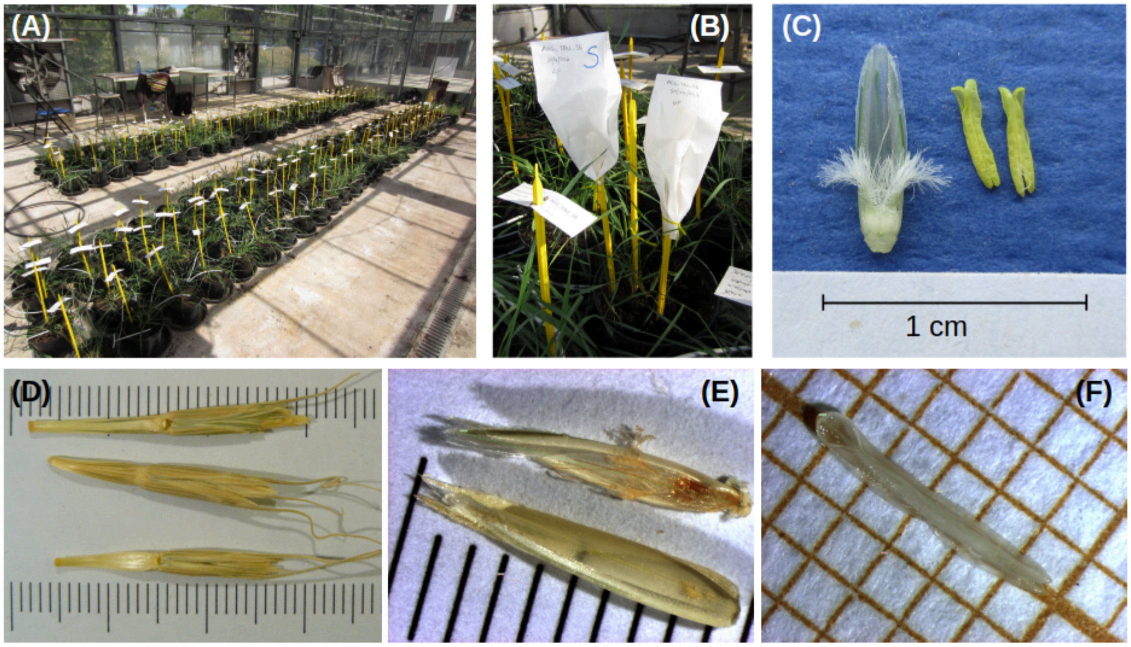


**Figure S12.** Experimental set up for the measure of the morphological traits in 16 species of Triticeae. Measures were taken with the software analySIS. A) Greenhouse set-up; B) Spikes bagged to prevent cross-fertilization; C) Ovary, stigma and anthers of *Ae. speltoides*; D) Three spikelets of *Ae. sharonensis*; E) Palea and lemna of a flower of *Ae. longissima*; F) Anther of *T. monococcum*.

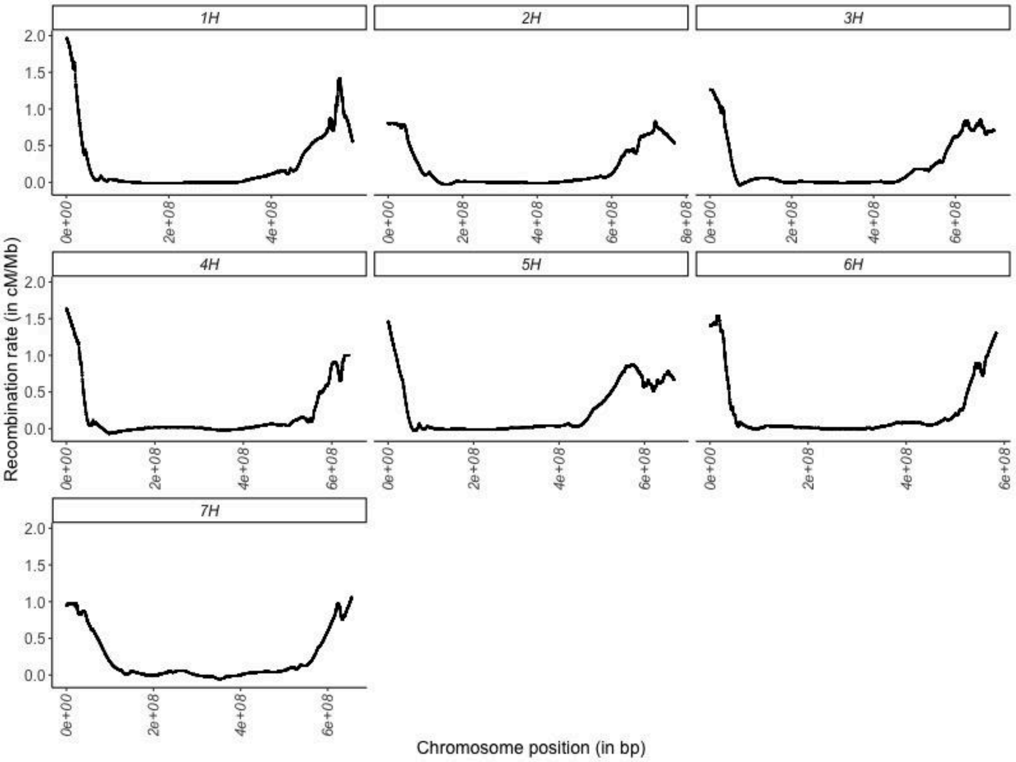
1H
2H
3H
4H
5H
6H
7H
0.0
0.5
1.0
1.5
2.0
0e+00
2e+08
4e+08
6e+08
8e+08
Recombination rate (in cM/Mb)
Chromosome position (in bp)

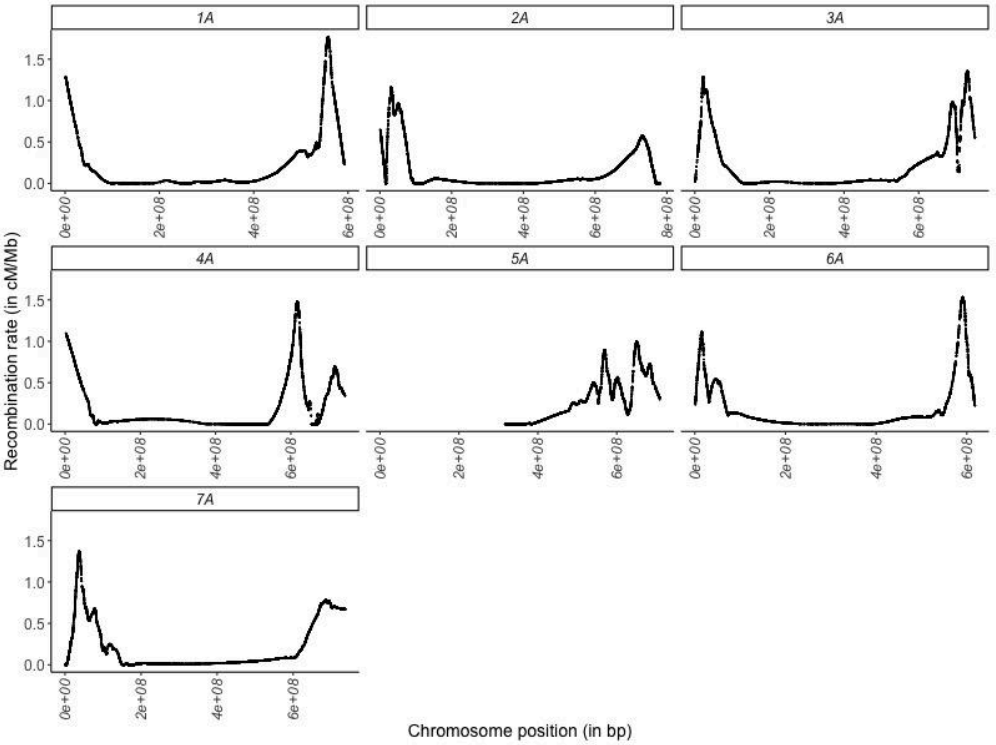
1A
2A
3A
4A
5A
6A
7A
0.0
0.5
1.0
1.5
0e+00
2e+08
4e+08
6e+08
8e+08
Recombination rate (in cM/Mb)
Chromosome position (in bp)

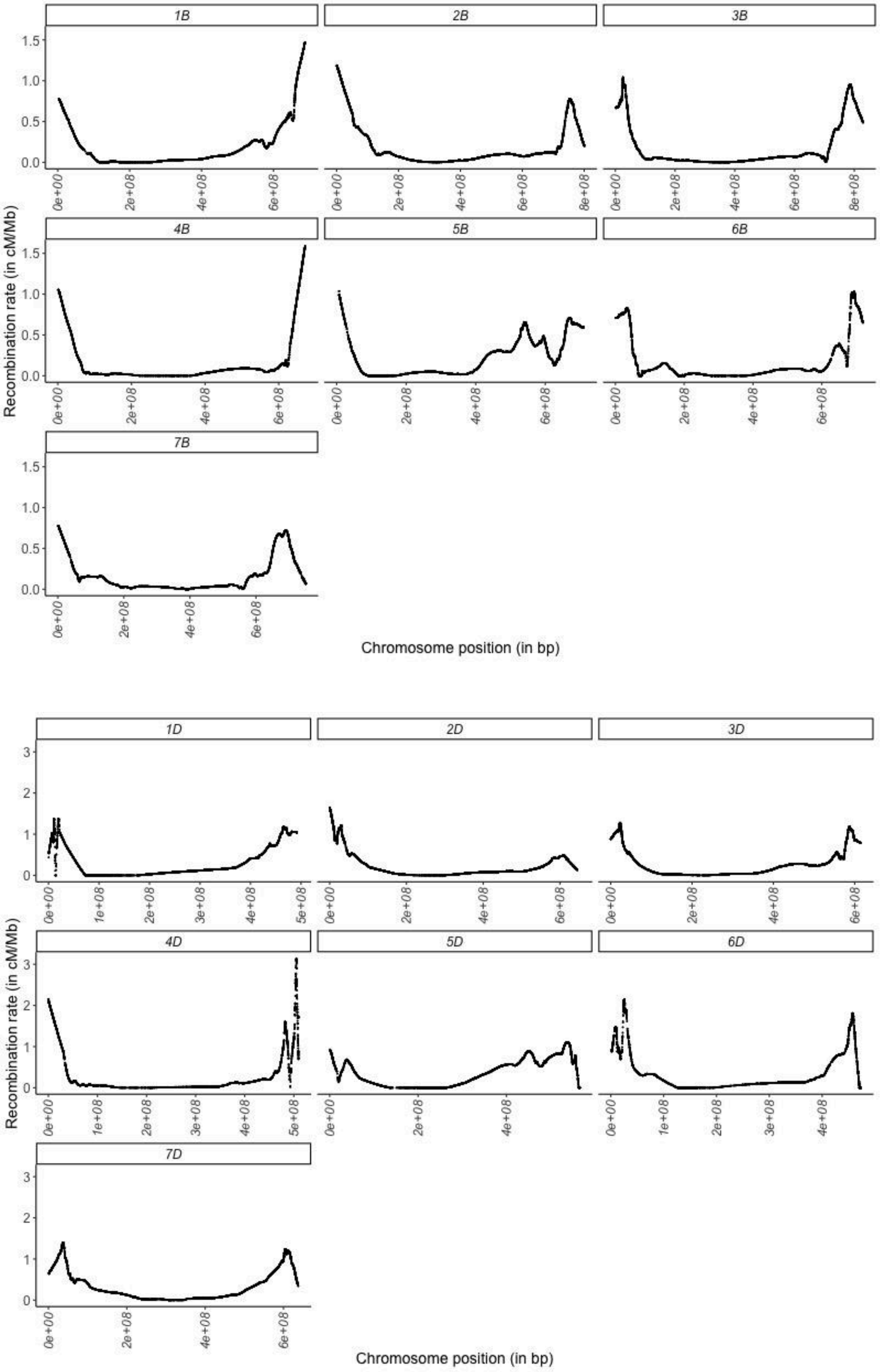


**Figure S13**: Recombination maps of the 7 chromosomes of barley (*Hordeum vulgare*) and of the three subgenomes of bread wheat (*Triticum aestivum*). The recombination maps were obtained as the local slope of the Marey map after smoothing using a loess function of degree two and a span parameter of 0.2.

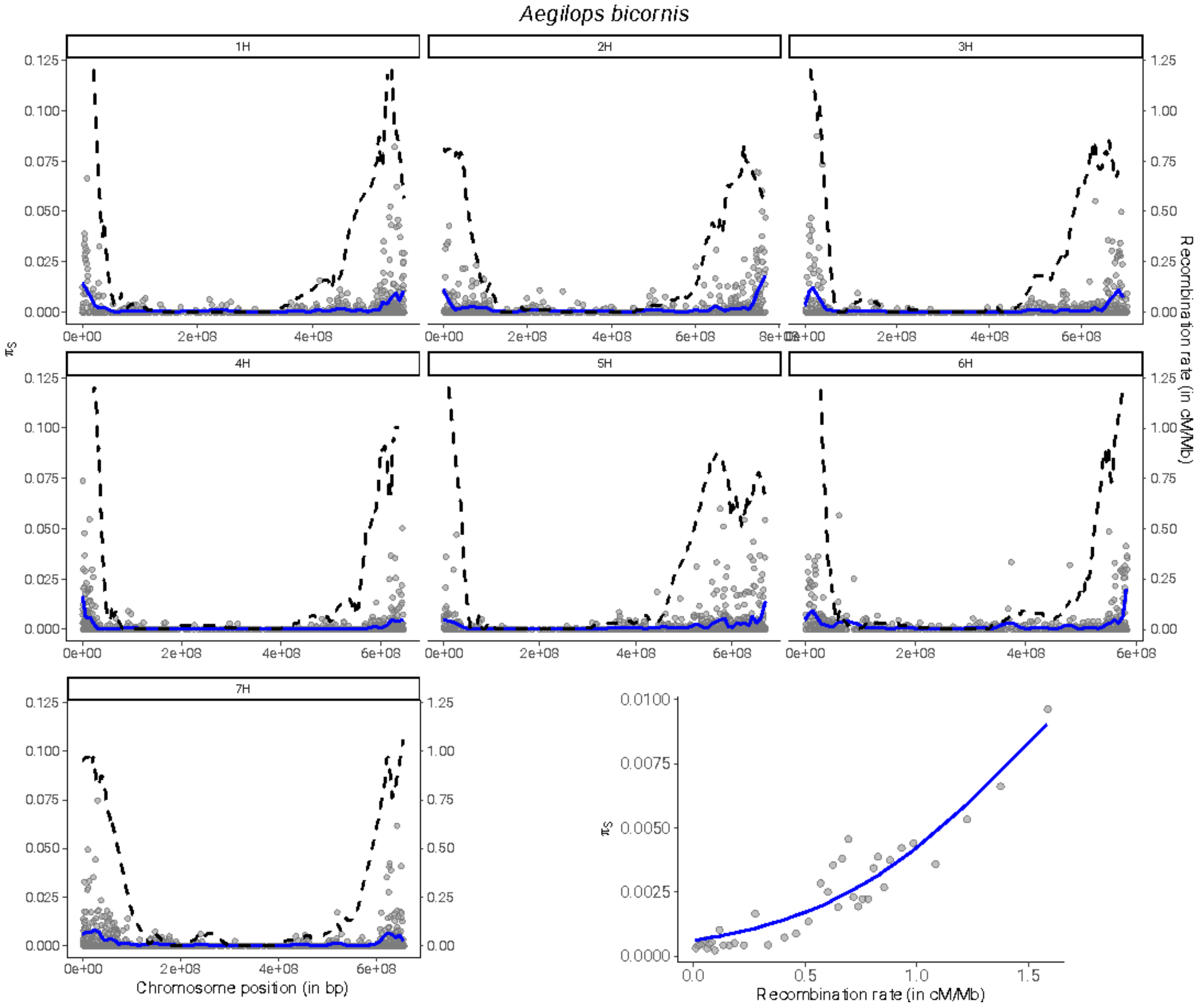
Aegilops bicornis
1H
2H
3H
4H
5H
6H
7H
πS
Recombination rate (in cM/Mb)
Chromosome position (in bp)
Recombination rate (in cM/Mb)

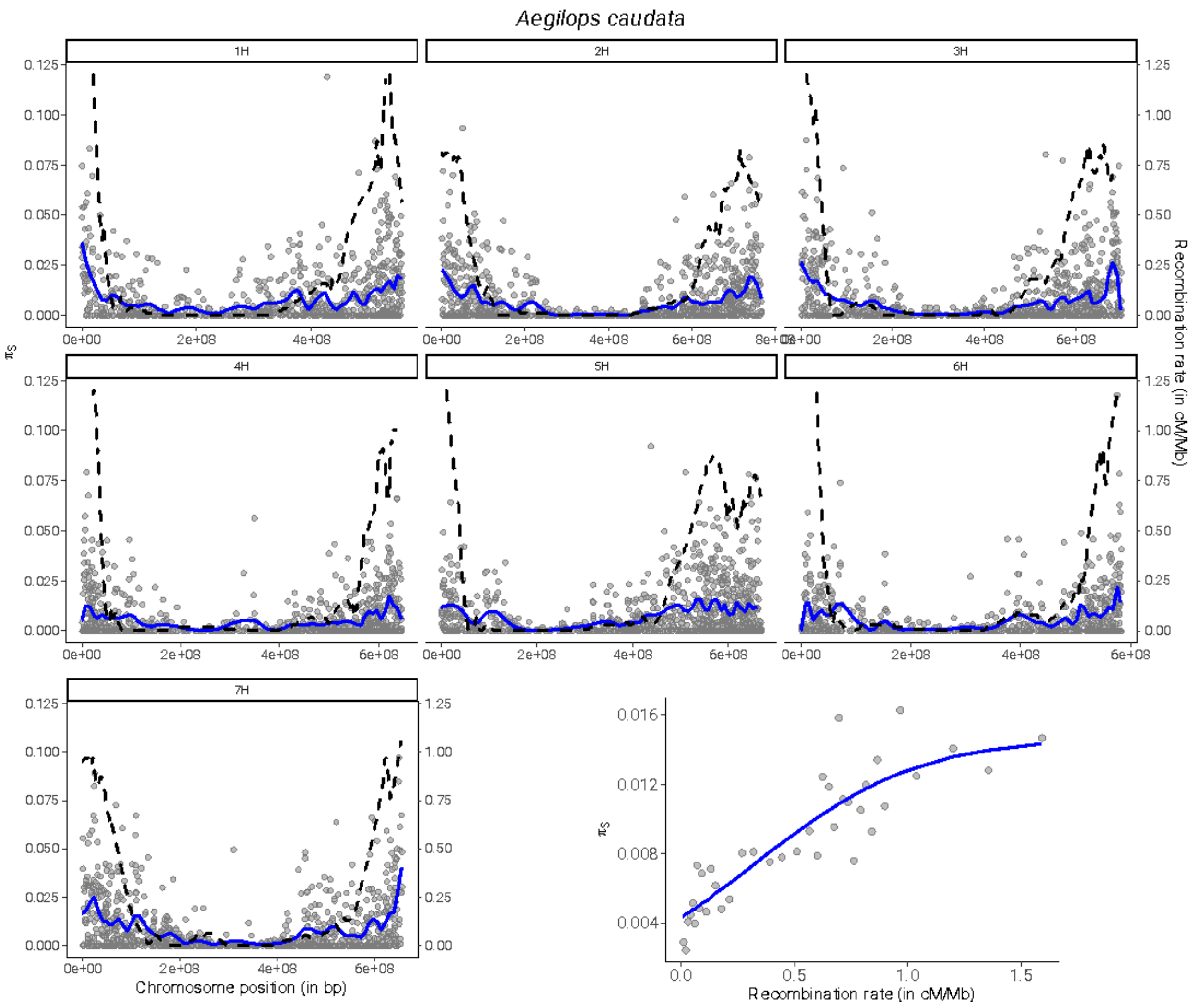
Aegilops caudata
1H
2H
3H
4H
5H
6H
7H
πS
Recombination rate (in cM/Mb)
Chromosome position (in bp)
0.125
0.100
0.075
0.050
0.025
0.000
1.25
1.00
0.75
0.50
0.25
0.00
0e+00
2e+08
4e+08
6e+08
8e+08
0.016
0.012
0.008
0.004
0.0
0.5
1.0
1.5
Recombination rate (in cM/Mb)

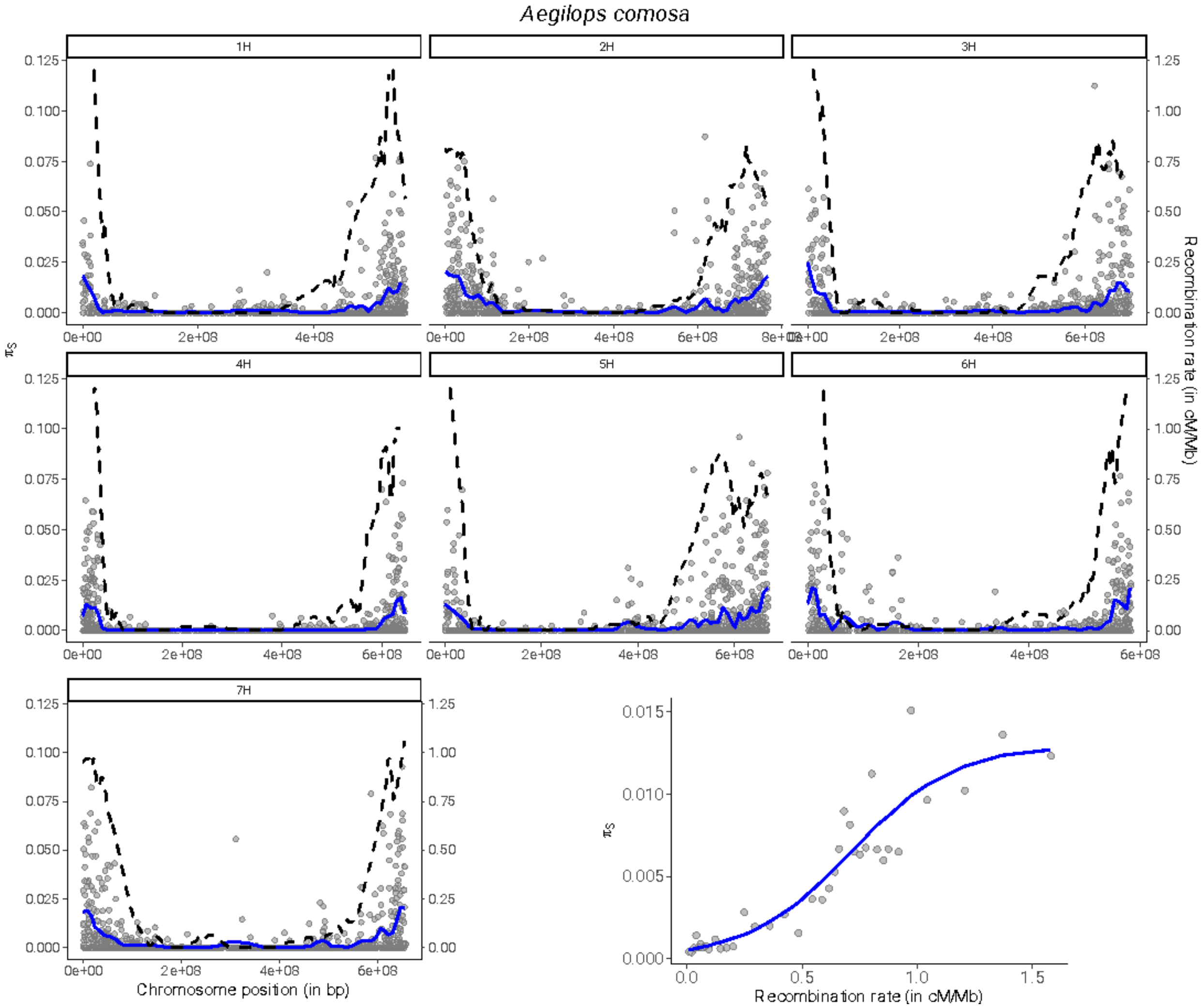
Aegilops comosa
1H
2H
3H
4H
5H
6H
7H
πS
Recombination rate (in cM/Mb)
Chromosome position (in bp)
Recombination rate (in cM/Mb)

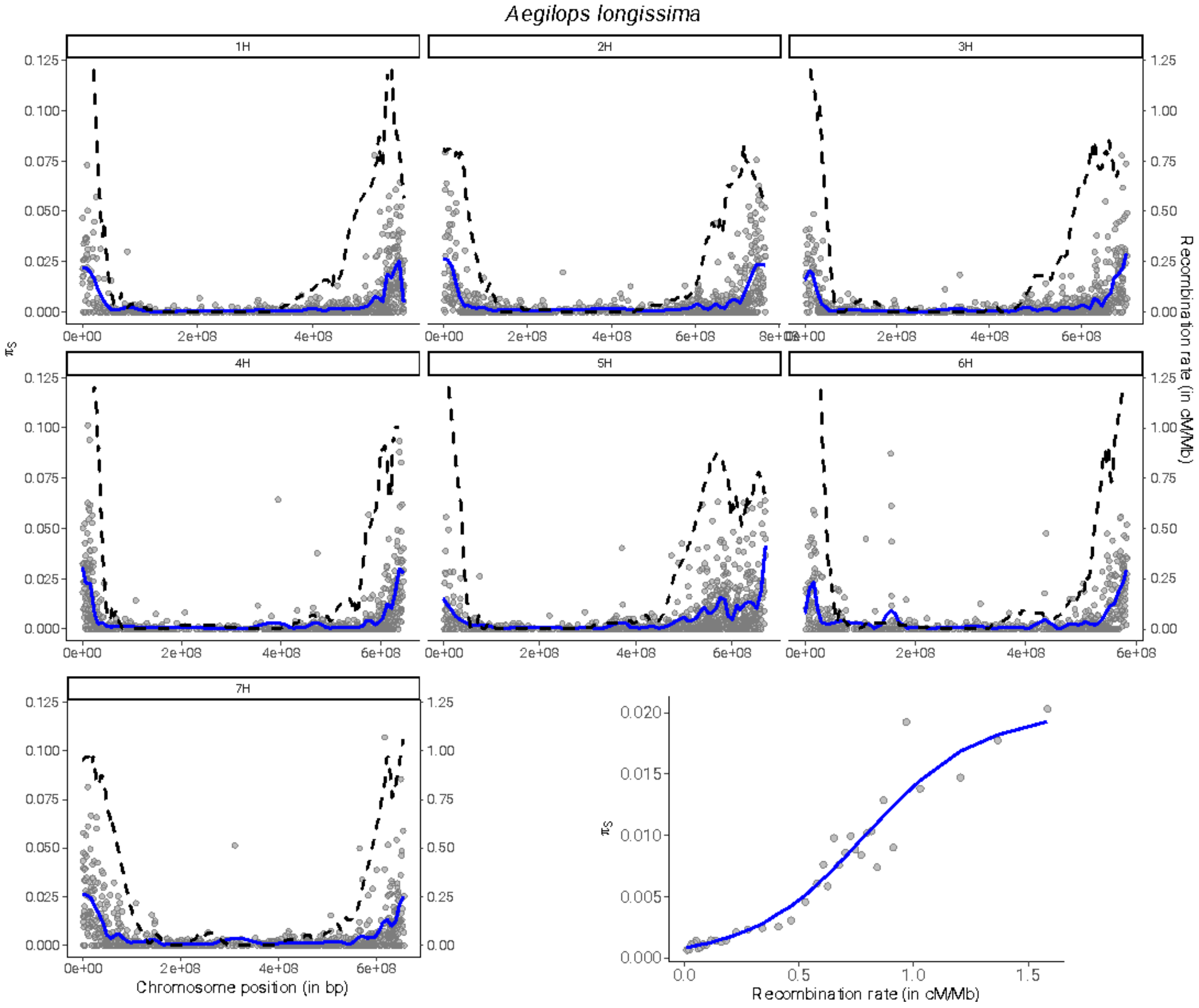

Aegilops longissima
1H
2H
3H
4H
5H
6H
7H
πS
Recombination rate (in cM/Mb)
Chromosome position (in bp)
Recombination rate (in cM/Mb)

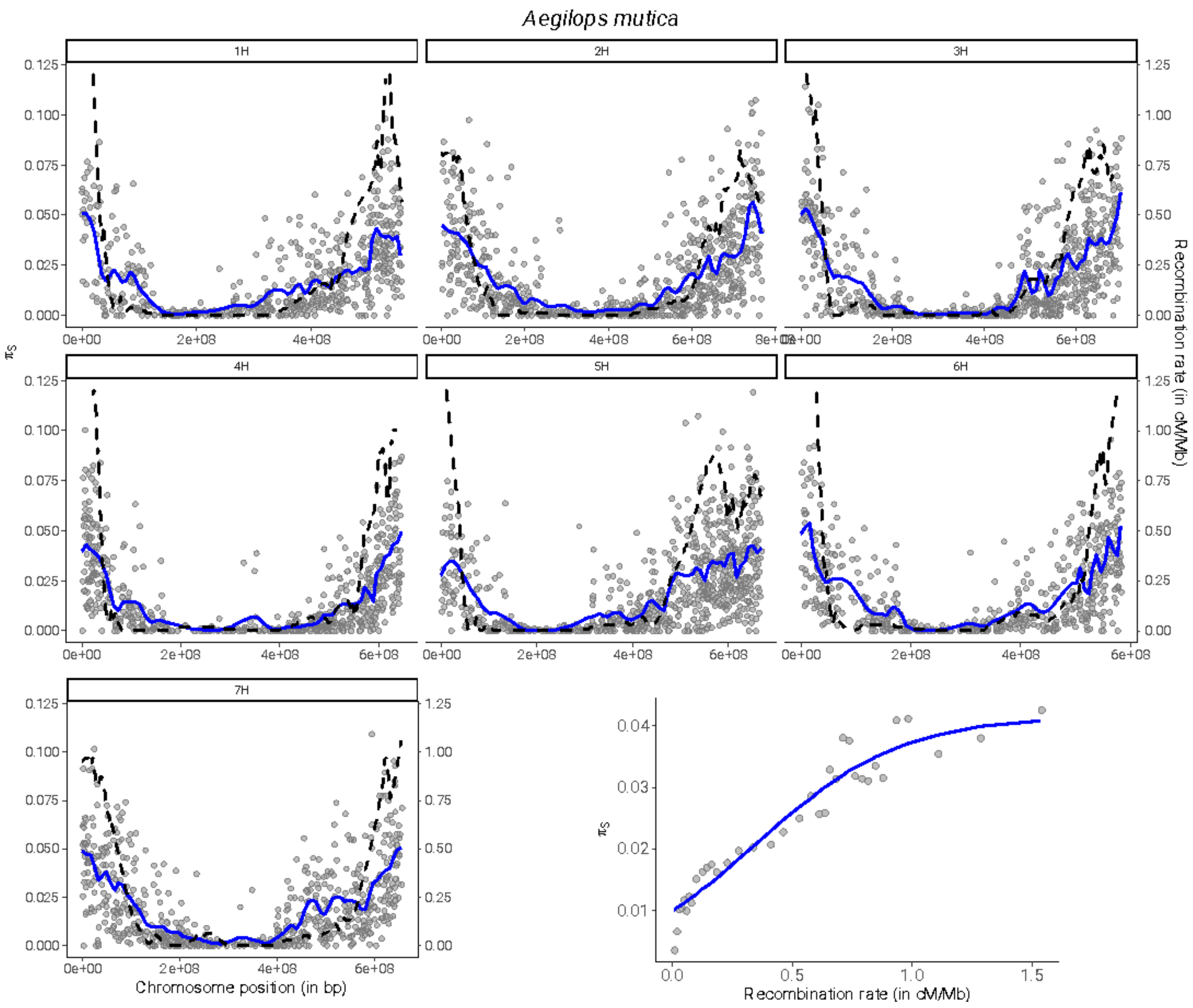

Aegilops mutica
1H
2H
3H
4H
5H
6H
7H
πS
Recombination rate (in cM/Mb)
Chromosome position (in bp)

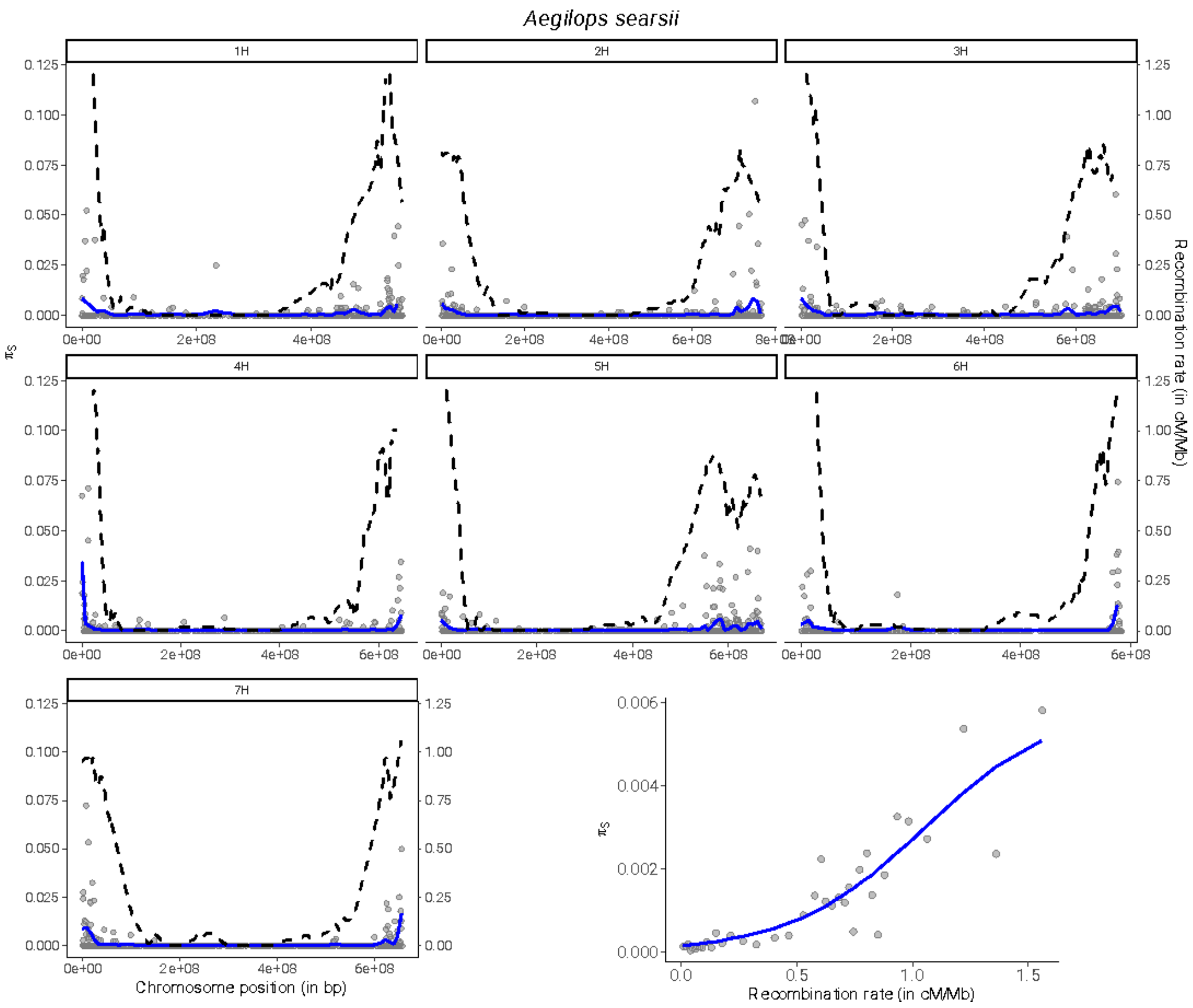

Aegilops searsii
1H
2H
3H
4H
5H
6H
7H
πS
Recombination rate (in cM/Mb)
Chromosome position (in bp)
Recombination rate (in cM/Mb)

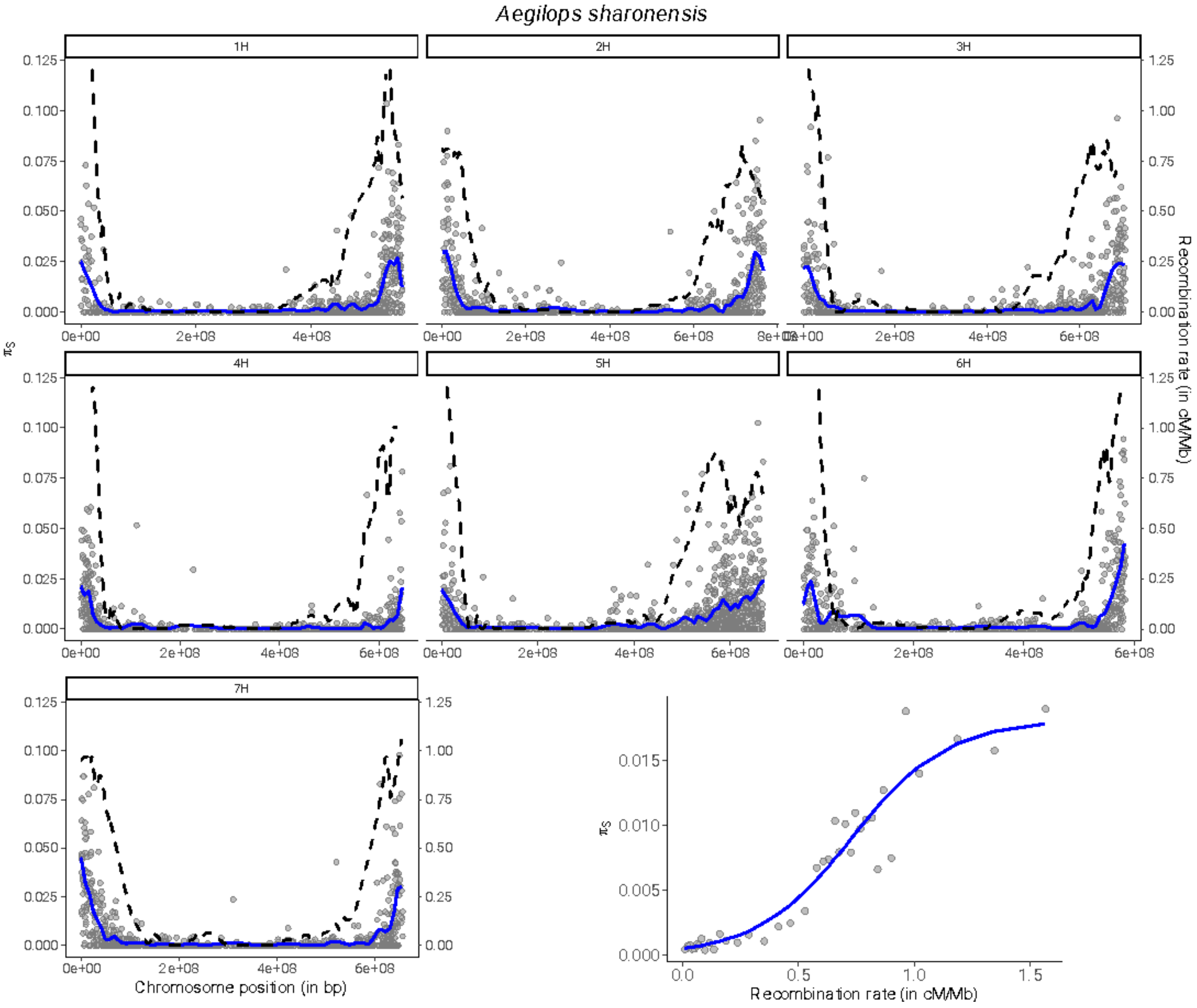
Aegilops sharonensis
1H
2H
3H
4H
5H
6H
7H
πS
Recombination rate (in cM/Mb)
Chromosome position (in bp)
0.125
0.100
0.075
0.050
0.025
0.000
1.25
1.00
0.75
0.50
0.25
0.00
0e+00
2e+08
4e+08
6e+08
8e+08
0.015
0.010
0.005
0.000
0.0
0.5
1.0
1.5

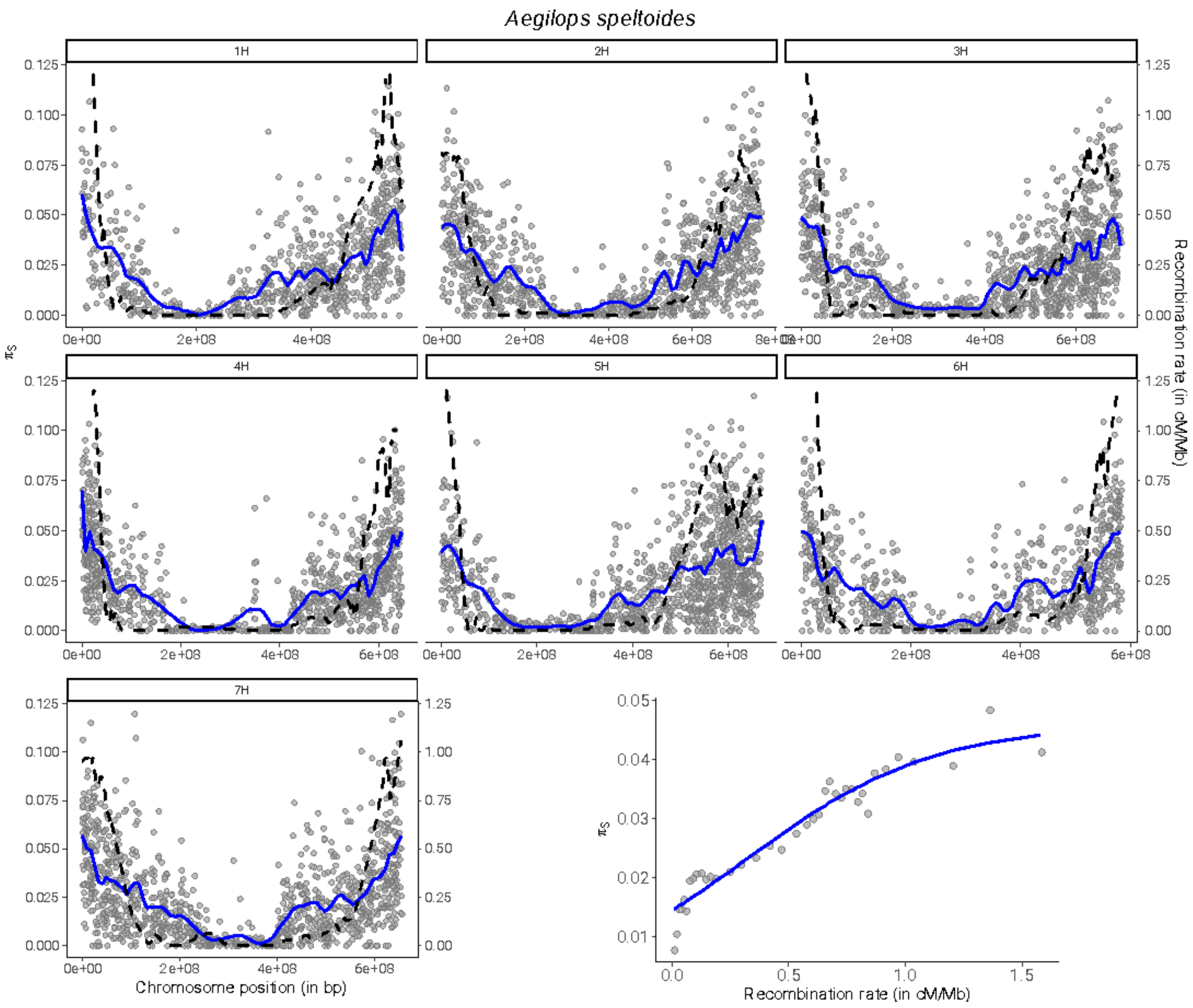

Aegilops speltoides
1H
2H
3H
4H
5H
6H
7H
πS
Recombination rate (in cM/Mb)
Chromosome position (in bp)
Recombination rate (in cM/Mb)

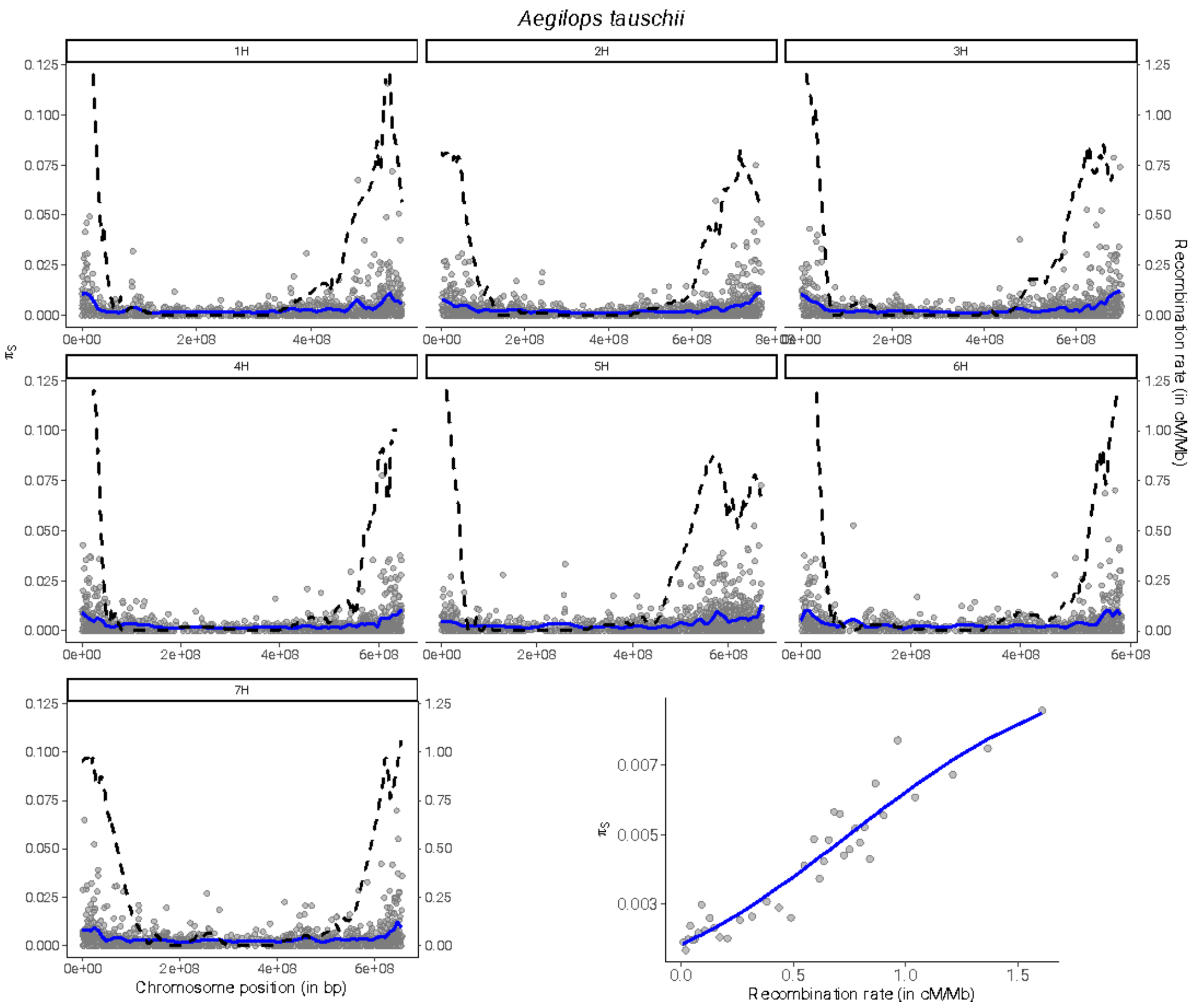

Aegilops tauschii
1H
2H
3H
4H
5H
6H
7H
πS
Recombination rate (in cM/Mb)
Chromosome position (in bp)

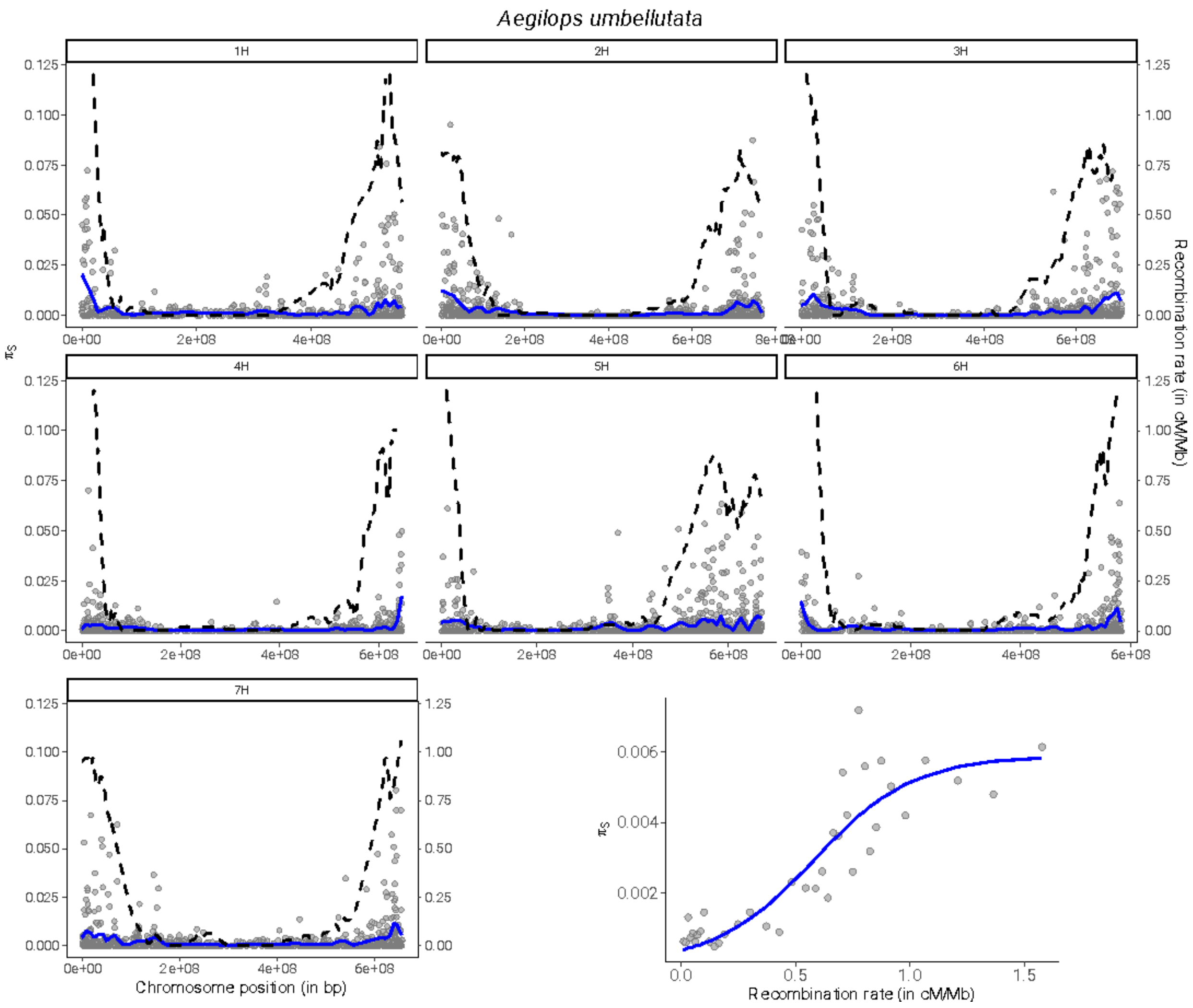

Aegilops umbellutata
1H
2H
3H
4H
5H
6H
7H
πS
Recombination rate (in cM/Mb)
Chromosome position (in bp)
Recombination rate (in cM/Mb)

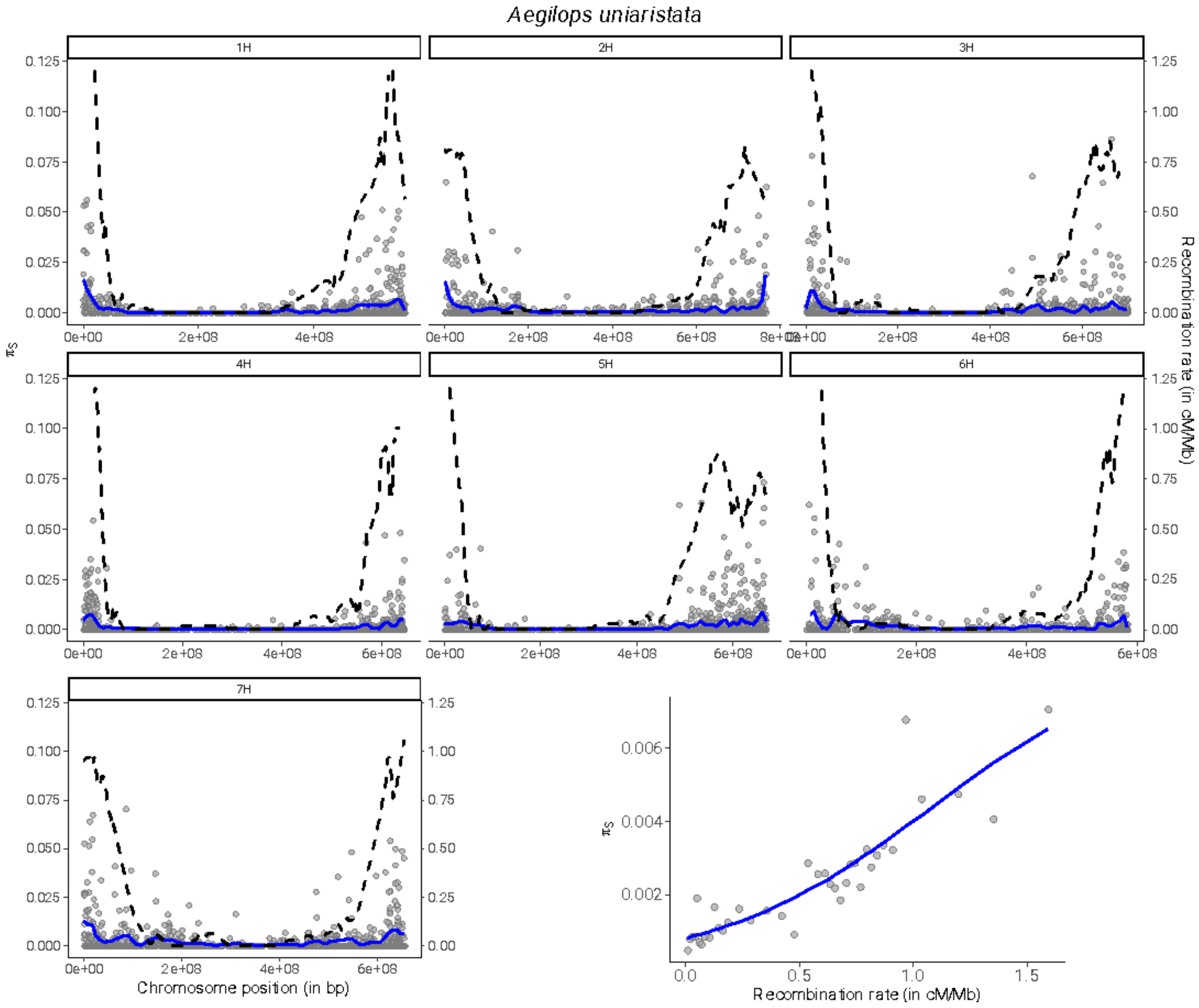

Aegilops uniaristata
1H
2H
3H
4H
5H
6H
7H
πS
Recombination rate (in cM/Mb)
Chromosome position (in bp)
Recombination rate (in cM/Mb)

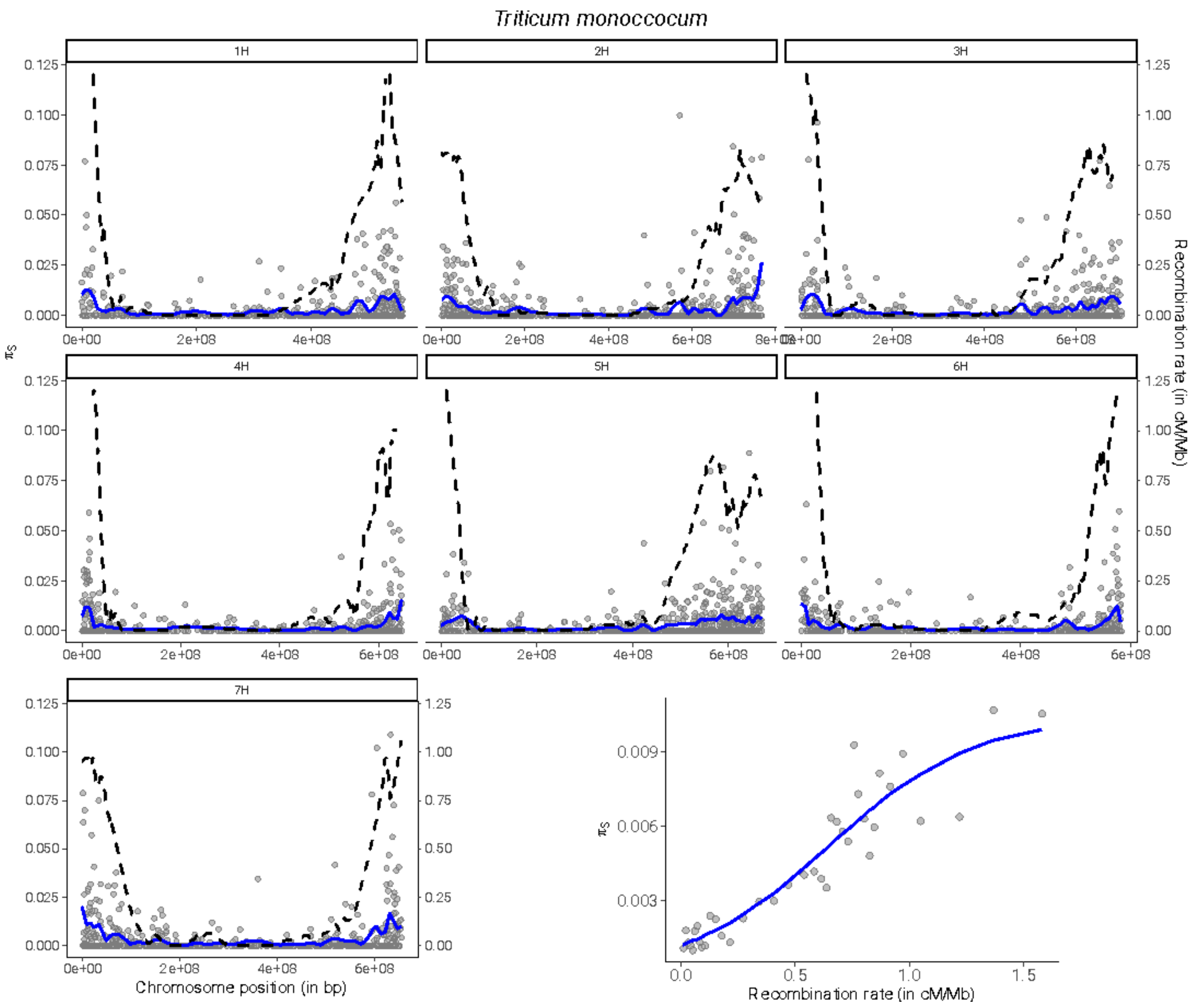
Triticum monoccocum
1H
2H
3H
4H
5H
6H
7H
πS
Recombination rate (in cM/Mb)
Chromosome position (in bp)
Recombination rate (in cM/Mb)

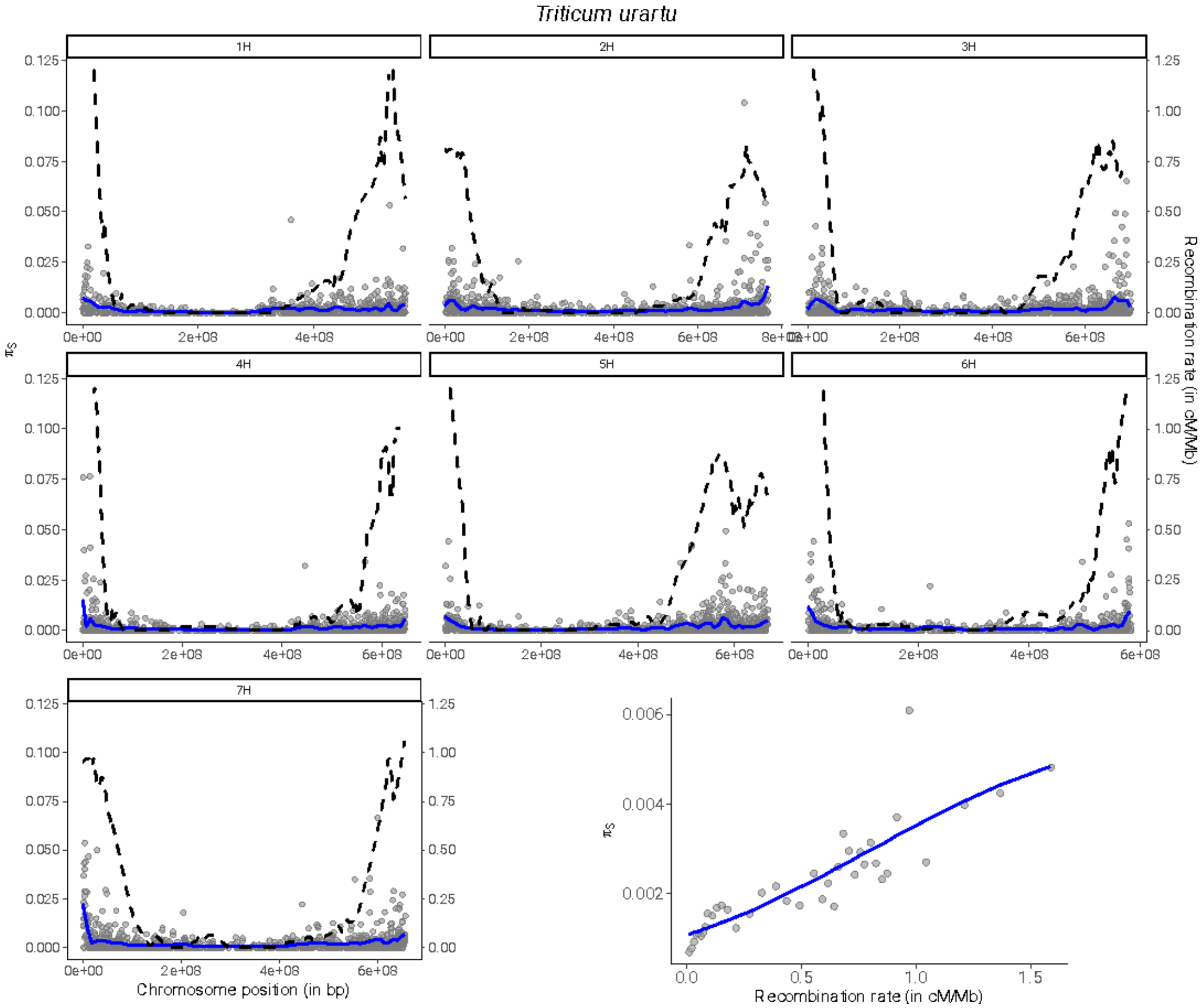


**Figure S14.** Mean synonymous diversity ($\pi_S$) along the seven chromosomes for all species (one figure per species). Each point corresponds to a contig mapped on the *Hordeum vulgare* genome. The blue line is the loess fitting function (degree=2, span=0.1). The dashed black line indicates *H. vulgare* recombination map (in cM/Mb) as in Fig. S12. The last bottom right figure is the mean $\pi_S$ computed in 50 quantiles of recombination. A sigmoid function is fitted (blue line).
